# microRNAs affecting development of body pigmentation in adult Drosophila melanogaster

**DOI:** 10.64898/2026.01.12.698815

**Authors:** Abigail M. Lamb, Jennifer A. Kennell, Eden W. McQueen, Evan J. Waldron, Erick X. Bayala, Patricia J. Wittkopp

## Abstract

Phenotypic development is regulated by multiple mechanisms that ensure tight control of gene expression. Post-transcriptional regulation, including the silencing or degradation of messenger RNAs by microRNAs (miRNAs), is an important component of this process. Here, we use gain-of-function and loss-of-function screens to examine the effects of miRNAs on cuticular pigmentation in adult *Drosophila melanogaster*. We found that 48 of 166 miRNAs ectopically expressed in a stripe along the dorsal side of developing flies were each sufficient to affect pigmentation. We also found that 22 of 41 miRNAs competitively inhibited in the same tissue visibly altered pigmentation, showing that they were necessary for adult pigmentation development. For each of the 15 miRNAs with opposing effects in the gain- and loss-of-function screens, computational tools identified possible targets among 93 genes previously reported to affect adult pigmentation. Using cell culture, we found that one of these miRNAs (*miR-8*) was able to regulate gene expression through 3’ UTR sequences from at least three pigmentation genes: *ebony*, *bric-a-brac 1*, and *bric-a-brac 2*. All three of these genes reduce development of black pigments, suggesting that *miR-8* coordinately regulates expression of multiple genes with similar effects on pigmentation. These data show that miRNAs are developmental regulators of body pigmentation, which could also allow them to contribute to pigmentation divergence, as has been shown for *miR-193* in butterflies.

## Introduction

Proper development of a multicellular organism requires strict control of gene expression, with genes expressed at the necessary time and place and in the proper amount and environmental context. This expression is controlled first by transcriptional regulation, in which *cis*-regulatory DNA sequences interact with transcription factors to determine when, where, and how much RNA is transcribed from a gene. After RNA transcripts are made, post-transcriptional regulation can further modify the expression of gene products by altering the stability, splicing, capping, polyadenylation, and translational efficiency of RNAs (Halbeisen et al. 2008). Small, non-coding RNAs called microRNAs (miRNAs) are important post-transcriptional regulators that work by guiding the RNA-induced silencing complex (RISC) to the 3’ UTRs of messenger RNA (mRNA) and preventing its translation into protein (Bartel 2018) (**Figure 1A**). Early studies of miRNAs in *Caenorhabditis elegans* showed that loss of a single miRNA’s function often had no discernable impacts on the phenotypes assayed (Miska et al. 2007), but more recent work has found that loss of miRNA function can have strong effects on many organismal traits, including viability (Chen et al. 2014; Fulga et al. 2015), lifespan (Chen et al. 2014), fertility (Chen et al. 2014), morphology (Arif et al. 2013; Fulga et al. 2015; Verma and Cohen 2015; Tian et al. 2024), and behavior (Chen et al. 2014; Picao-Osorio et al. 2015; Picao-Osorio et al. 2017; Garaulet et al. 2020).

**Figure 1.**
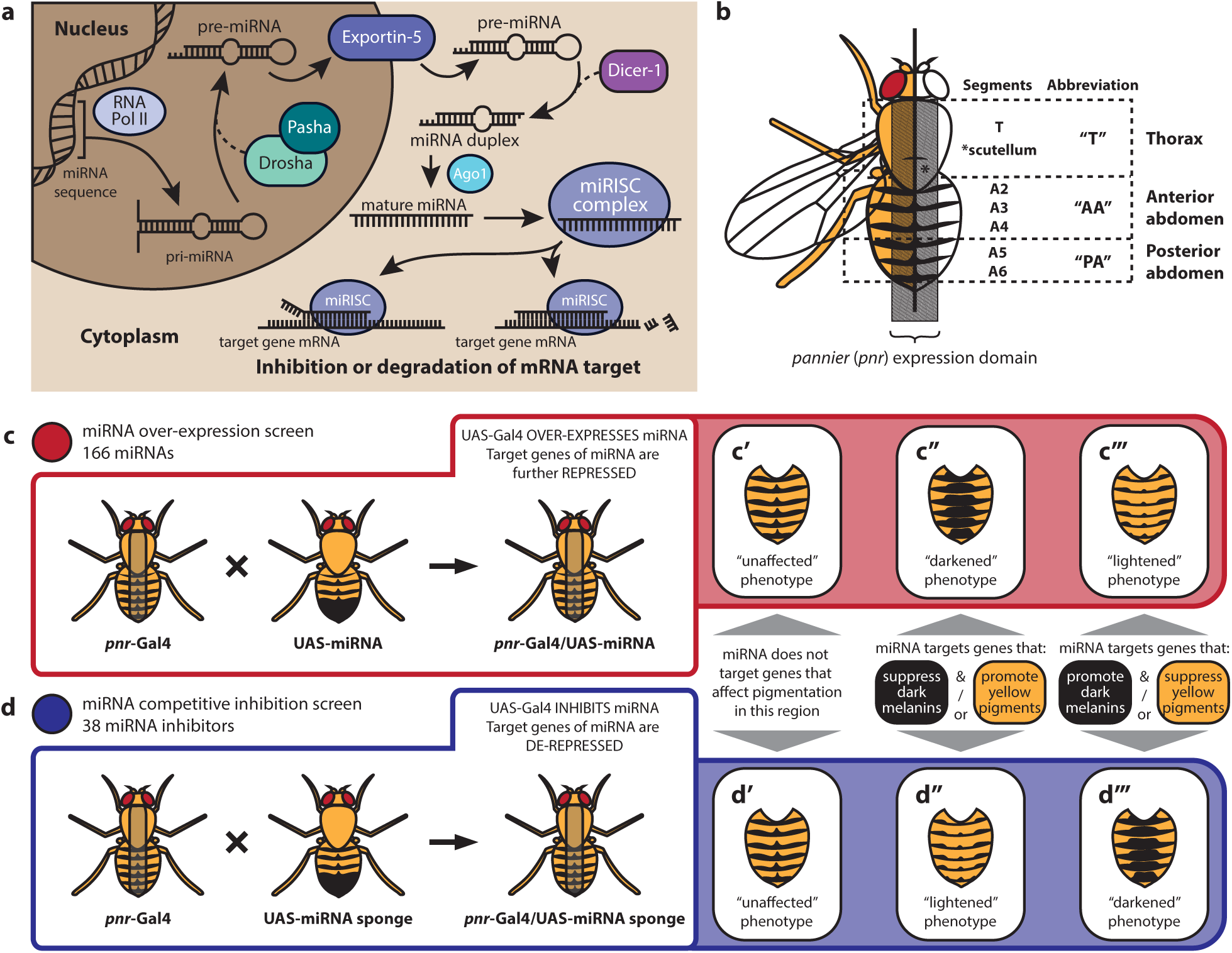
Overview of miRNA overexpression and competitive inhibition screens. **(A)** Schematic shows canonical miRNA biogenesis and mechanism of post-transcriptional repression. RNA polymerase II (Pol II) transcribes the pri-miRNA. Pri-miRNA transcript is cleaved by the microprocessor complex, composed of *Drosha* and *Pasha* (*DGCR8*), producing the precursor-miRNA (pre-miRNA). The pre-miRNA is exported to the cytoplasm by *Exportin5*, where it associates with the RISC loading complex (miRLC), which consists of *Dicer*, *TAR RNA binding protein* (*TRBP*), and *Argonaute 2* (*Ago2*). The pre-miRNA is processed by *Dicer* to produce the miRNA duplex. *Ago2* unwinds the duplex and maintains association with the mature miRNA, forming the miRNA-induced silencing complex (miRISC). In the presence of the target mRNA, the miRISC binds to the target’s 3′ untranslated region, resulting in translational repression or degradation. **(B)** Model of a female fly showing the location of the *pannier* expression domain in a region flanking the dorsal midline, as well as the three anatomical regions assessed for pigmentation: thorax (T), anterior abdomen (AA) and posterior abdomen (PA). The thorax region includes the scutellum, anterior abdomen refers to abdominal segments A2, A3 and A4, and posterior abdomen refers to abdominal segments A5 and A6. **(C, D)** Screening for miRNAs affecting abdominal pigmentation: Females carrying the *pnr-Gal4* driver were crossed to males carrying a UAS-miRNA transgene for the overexpression screen (**C**), or to males carrying a UAS-miRNA sponge for the competitive inhibition screen (**D**). Progeny of these crosses have altered miRNA in the *pnr-Gal4* expression domain and were scored as having “unaffected”, “darkened”, or “lightened” pigmentation based on a visual comparison of pigmentation in the *pnr-Gal4* expression domain to the cuticle flanking that domain. Schematics show these three categories of effects and what they suggest about the function of the miRNA **C’/D’:** The miRNA does not target genes that actually or potentially affect pigmentation in the dorsal midline. **C’’/D’’:** The miRNA targets genes that actually or potentially suppress dark melanins or promote yellow pigmentation in the dorsal midline. **C’’’/D’’’:** The miRNA targets genes that actually or potentially promote dark melanins or suppress yellow pigmentation in the dorsal midline.

Here, we advance our understanding of miRNA-mediated gene regulation affecting a complex trait by investigating the role of miRNAs in the development of body pigmentation in adult *Drosophila melanogaster*. This trait has been used as a model to study the regulation of gene expression, developmental processes, and mechanisms of phenotypic evolution (Massey and Wittkopp 2016; Rebeiz and Williams 2017). Prior work has identified many genes involved in the synthesis of pigments that comprise adult pigmentation patterns (Massey and Wittkopp 2016) as well as transcription factors that regulate (either directly or indirectly) the expression of those genes (Rogers et al. 2014; Kalay et al. 2016). Previous studies have also identified three miRNAs affecting adult pigmentation in *D. melanogaster: miR-8* and *miR-193* have been shown to promote darker pigmentation in the abdomen (Kennell et al. 2012; Tian et al. 2024) and *miR-33* has been shown to repress this dark pigmentation in the abdomen (Clerbaux et al. 2021). However, the full extent to which miRNAs regulate pigmentation development in *D. melanogaster*, and whether miRNAs interact directly with pigmentation genes, remains unknown.

To determine whether additional miRNAs affect development of adult pigment patterns in *D. melanogaster*, we performed gain-of-function and loss-of-function genetic screens. First, we used *pnr-Gal4* to express each of 166 miRNAs in a stripe along the dorsal side of developing flies, finding that 29% of the miRNAs tested (48/166) were sufficient to cause a visible change in pigmentation of the adult cuticle. Next, we used competitive inhibition to reduce activity of 41 miRNAs, finding that 22 were also necessary for the development of normal pigmentation. For 15 of these 22 miRNAs, the phenotype resulting from reducing its availability was the opposite of the phenotype resulting from increasing it, suggesting that these 15 miRNAs play roles in *D. melanogaster* pigmentation development. Using the TargetScan database (Agarwal et al. 2018), we found that these miRNAs have the potential to regulate expression of many genes previously shown to impact adult pigmentation in *D. melanogaster.* Cell culture experiments confirmed that one of these miRNAs (*miR-8*) was able to repress expression of a reporter gene containing 3’ UTR sequences from several pigmentation genes that promote lighter pigmentation. It remains to be seen whether *mir-8* directly regulates these genes *in vivo* and how the other miRNAs identified here exert their effects on pigmentation. Nevertheless, this study, along with work on other traits (e.g., Tsang et al. 2010; Barca-Mayo and De Pietri Tonelli 2014; Ben-Hamo and Efroni 2015; Lui et al. 2015), suggests that miRNAs influence trait development by coordinately regulating expression of multiple genes with related functions.

## Materials and Methods

### Fly strains and crosses

UAS-miRNA lines were obtained from the FlyORF Zurich ORFeome Project miRNA collection (www.flyORF.ch, (Schertel et al. 2012). For the overexpression screen, two virgin female *pnr*-Gal4/TM6B flies per vial were crossed to male UAS-miRNA flies for each miRNA tested (crosses were set for 172 total UAS-miRNA lines in 8 batches conducted across ∼10 weeks). miRNA overexpression crosses were reared in a 23^°^C incubator on a 12-hour light/dark cycle. For the competitive inhibition screen, two virgin females from the same *pnr*-Gal4 line were crossed to each UAS-miRNA sponge line (Bloomington Drosophila Stock Center, Fulga et al. 2015). Stock information for all UAS-miRNA and UAS-miRNA sponge lines is listed in **Supplementary Table S1**.

Because we anticipated potentially smaller effects from competitive inhibition than from overexpression, we set competitive inhibition crosses at two temperature extremes (see main text). We reasoned that the more darkly pigmented flies reared at 18^°^C would allow greater visibility of lightening effects from miRNA inhibition, while lighter flies reared at 28^°^C would allow greater visibility of darkening effects. All flies were aged 3-5 days after eclosion before phenotyping. Each fly’s pigmentation outside the *pnr* expression domain was used as an internal control for comparison to the effects of miRNA overexpression in the dorsal midline.

UAS-*ebony* (Wittkopp et al. 2002) and UAS-*ebony* RNAi (BDSC_28612; Perkins et al. 2015) were crossed to *pnr*-Gal4 as positive controls for lightened and darkened midline pigmentation, respectively. Each fly was observed by A. L., at which time the eclosion date, collection date, genotype, sex, and phenotypes were recorded in a spreadsheet for each screen (**Supplementary Table S2, S3**). Flies with mutant phenotypes consistent with balancer chromosomes were documented with the identity of the balancer chromosome. Flies inheriting both *pnr*-Gal4 and a UAS transgene were documented categorically as unaffected, lightened, darkened, or both lightened and darkened, with a short description of the phenotype, segment(s) affected, and any conspicuous developmental defects. Additional crosses were set to compare *pnr*-Gal4/UAS-miRNA flies to a genetically similar negative control by first crossing the *pnr*-Gal4 to a double balancer line (w- ;Sco/CyO;MKRS/TM6B, provided by Scott Pletcher, University of Michigan) to obtain *pnr*-Gal4/MKRS flies. Only flies carrying the MKRS balancer chromosome were used because the TM6B balancer chromosome contains a loss-of-function mutant *ebony* allele that visibly affects pigmentation. We crossed *pnr*-Gal4/MKRS flies to flies carrying the UAS-*miR-8*, UAS-*miR-92b*, UAS-*miR-33*, and UAS-*miR-279* transgenes, rearing and collecting flies in the same conditions and manner as for the UAS-miRNA screen crosses described above.

Strains carrying knockout alleles of *miR-92b* (stock #58938, w[*]; TI{w[+mW.hs]=TI}mir-92b[KO]/TM3, Sb[1]), *miR-193* (stock #58898, w[*]; TI{w[+mW.hs]=GAL4}mir-193[KO]/TM3, P{w[+mC]=GAL4-twi.G}2.3, P{UAS-2xEGFP}AH2.3, Sb[1] Ser[1]), and *miR-276a* (stock #58906, w[*]; TI{w[+mW.hs]=TI}mir-276a[KO]/TM3, P{w[+mC]=GAL4-twi.G}2.3, P{UAS-2xEGFP}AH2.3, Sb[1] Ser[1]) were obtained from the Bloomington Drosophila Stock Center.

### Sample preparation and imaging

Flies displaying lightened or darkened pigmentation phenotypes were placed in a solution of 10% glycerol in ethanol and labeled with the associated cross information. Flies from two randomly chosen *pnr*-Gal4/UAS-miRNA crosses for which all individuals were classified as displaying an “unaffected” phenotype (UAS-*miR-100* and UAS-*miR-1014*) were also stored in 10% glycerol in ethanol for comparison purposes. A subset of flies with notable phenotypes (including those shown in **Figure 2)** were photographed immediately after collection and before storage, using a Leica MZ6 stereomicroscope equipped with a ring light and Scion (CFW-1308C) camera operated via TWAIN driver in Adobe Photoshop. Dorsal abdominal cuticles were dissected from the offspring of the *pnr*-Gal4/MKRS x UAS-miRNA crosses described in “Fly strains and crosses” for **Figure 4** and **Figure 6B**. Dissection and mounting were performed as described in John et al. (2016) before imaging. For the images in Figures 4, flies preserved from the screen experiments were partially embedded in 1% agar in white centrifuge tube caps, which were then flooded with 95% ethanol before imaging. All photographs in **Figure 4** and **Figure 6B** were taken using the same equipment described above.

**Figure 2.**
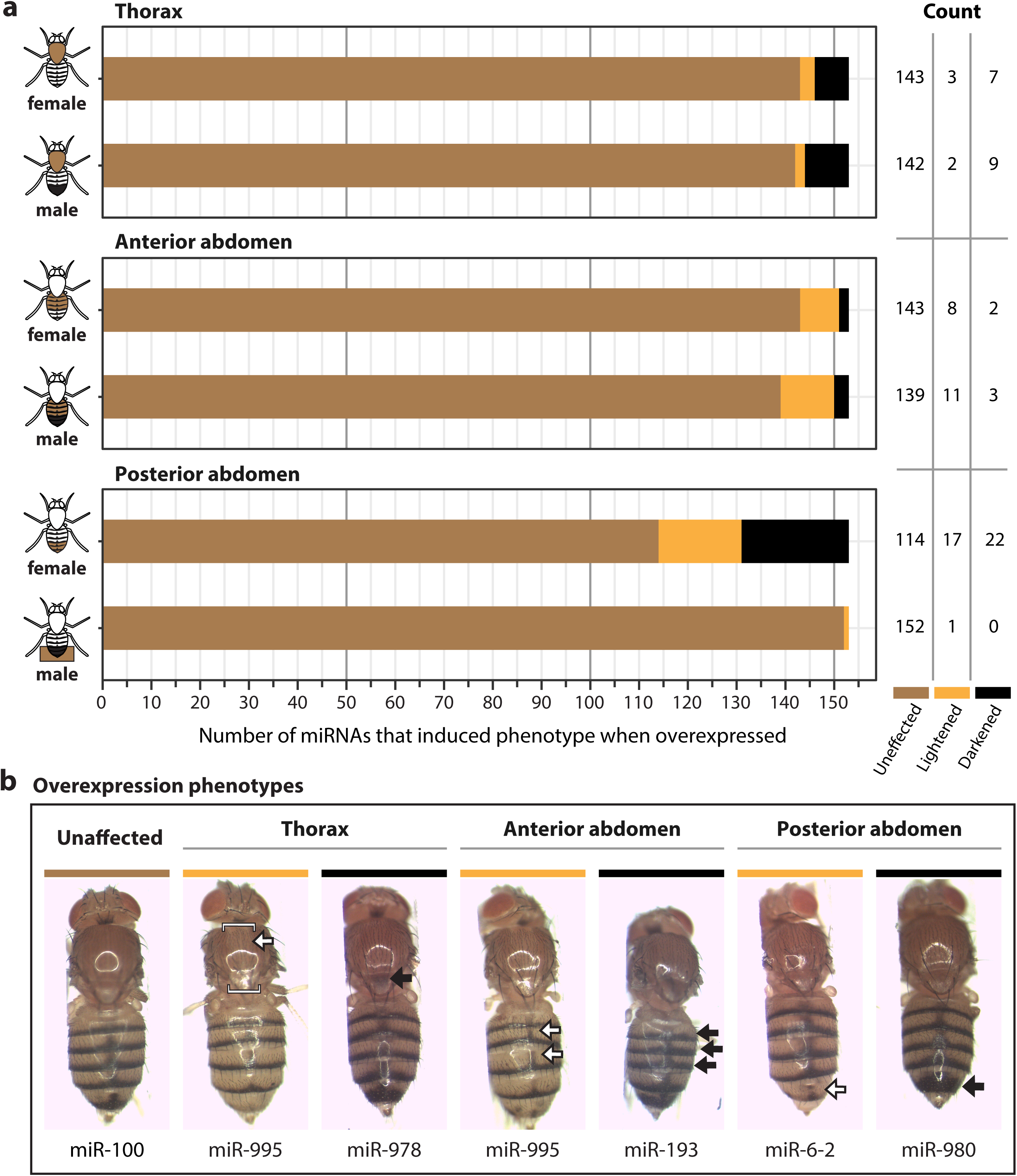
Pigmentation phenotypes resulting from overexpressing each of 153 miRNAs. **(A)** The relative frequencies of miRNAs that had no observable effect on pigmentation (“Unaffected”) or were sufficient to lighten (“Lightened”) or darken (“Darkened”) pigmentation are shown for both male and female flies. Each row shows data from one of the three body regions scored along the anterior-posterior body axis: thorax, anterior abdomen (segments A2-A4), and posterior abdomen (segments A5-A6). **(B)** Images show examples of flies classified as having “Unaffected”, “Lightened”, or “Darkened” pigmentation in the *pnr-Gal4* expression domain relative to the flanking regions. Black arrows indicate regions with darkened pigmentation in the *pnr-Gal4* expression domain, and white arrows indicate regions with lightened pigmentation in the *pnr-Gal4* expression domain. A fly classified as having pigmentation unaffected by altered miRNA expression is also shown. In each case, the miRNA overexpressed in the *pnr-Gal4* expression domain is shown.

### Analysis of predicted miRNA target genes

We compiled a list of 93 genes with experimentally validated effects on pigmentation by searching literature and annotations from flybase.org (Thurmond et al. 2019). First, we collected the names and phenotypes associated with all genes described in (a) three reviews on Drosophila pigmentation (Wright 1987; Wittkopp et al. 2003; Massey and Wittkopp 2016), (b) two large-scale transcription factor RNAi screens for pigmentation regulators (Rogers et al. 2014; Kalay et al. 2016), and (c) a genome-wide association study (GWAS) of within species variation for body pigmentation using the Drosophila Genome Reference Panel (Dembeck, Huang, Magwire, et al. 2015). We then searched Gene Ontology annotations for “Biological Process” on Flybase.org, filtering for the following terms: “negative regulation of developmental pigmentation”, “positive regulation of developmental pigmentation”, “regulation of cuticle pigmentation”, “regulation of eye pigmentation”, “regulation of female pigmentation”, “regulation of male pigmentation”, “regulation of pigment cell differentiation”, “regulation of adult chitin-containing cuticle pigmentation”. We then checked the references for each gene identified by this search, excluded all that were solely associated with pigmentation in structures other than adult cuticle, and reviewed the evidence supporting any remaining genes that were not described in the previously described sources. We only included genes where the supporting data presented in the referenced studies included descriptions or images of mutant or RNAi phenotypes associated with the gene in question. The results of this work are shown in **Supplementary Table S4**.

To predict potentially direct miRNA-mRNA regulatory interactions, we downloaded all predictions for genome wide matches to both “conserved” and “nonconserved” miRNA seed sites in the 3’ UTRs of the most highly expressed transcript of each gene in the *D. melanogaster* genome from TargetScanFly v7.2 (http://www.targetscan.org/fly_72/, Agarwal et al. 2018). We then filtered this database to only include the 3’ UTRs of genes listed in **Supplementary Table S4** and the 14 miRNA seeds associated with miRNAs that produced opposite phenotypes when overexpressed versus competitively inhibited in the screens described in this study (**Table 2**).

### Cell culture assays of predicted miR-8 target genes

To overexpress *miR-8* in cultured *Drosophila* cells, the *miR-8* gene was cloned into a pAc5.1-V5/His-a expression vector (Invitrogen) containing the strong, constitutively active promoter from the *Drosophila* actin 5C gene (pAc-miR-8). To test for binding to the 3’ UTR sequences from putative target genes, the 3’ UTR sequences were cloned into a pAc-LacZ vector (Invitrogen) containing an upstream in-frame stop codon, a polyadenylation signal, and the same actin 5C promoter using the restriction enzymes NotI and XhoI (pAc-LacZ-3’UTR constructs). The full-length 3’ UTR was included in the construct for *ebony* (203 bp) For *bab1*, which has a 1953 bp 3’UTR, we used a 683 bp region including the predicted *miR-8* binding site. Similarly, for *bab2*, which has a 1539 bp 3’ UTR, we used a 529bp region including the predicted *miR-8* binding site. To disrupt the predicted *miR-8* binding sites in these *ebony*, *bab1* and *bab2* reporter genes, we mutated the 3’ UTR sequence complementary to the 7mer *miR-8* seed sequence from AGTATTA to CTGCGGC. A negative control reporter gene lacking any 3’UTR between the stop codon of LacZ and the polyadenylation signal, as well as a positive control reporter gene containing two perfect complements to the mature *miR-8* sequence, were also analyzed. A plasmid containing the luciferase gene under the control of the actin promoter was used as a control for transfection efficiency.

For each experiment, a LacZ reporter gene, the luciferase control, and an expression plasmid that did or did not express *miR-8* were transfected into Kc167 cells using the protocol described in Blauwkamp et al. (2008). Briefly, we plated 500uL of cells in 24-well plates at a density of 1x10^6^ cells/mL and allowed these cell cultures to grow for 24 hours at 25°C before transfection. Cells were transiently transfected using FuGENE 6 transfection reagent (Promega) with a total of 150ng of plasmid DNA (125ng of either pAc-*miR-8* or empty pAC vector, 12.5ng of a LacZ reporter, and 12.5ng of pAc-Luciferase). Approximately 36 hours after transfection, cells were collected, lysed, and activity of Beta-galactosidase and Luciferase was measured using GalactoStar (Invitrogen) and Tropix LucScreen (Applied Biosystems). All transfections were performed in duplicate on at least 3 days in the Kennell lab. For each reporter gene, these data were analyzed by fitting Beta-galactosidase activity to a linear model including Luciferase activity as a covariate and both experiment date and *miR-8* expression (presence or absence) as main effects. Estimated marginal means (also called least-squares means or LSmeans) for Beta-galactosidase activity and their 95% confidence intervals were calculated from this model using the *emmeans* R package and normalized by dividing by the LSmean from the sample without overexpression of *miR-8*. Pairwise contrasts (t-test) were performed for each reporter gene between the LSmeans of samples that did and did not express *miR-8*.

All of these LacZ reporter gene constructs except the mutant version of *bab2*, plus LacZ reporters containing the full 1953 bp *bab1* 3’ UTR sequence and 3’ UTR sequences from *black* (268 bp), *homothorax* (*hth*) (3816 bp), *pale* (1614 bp), and *Hormone receptor-like in 38* (*Hr38*) (3348 bp), were also tested for activity with and without overexpression of *miR-8* in *Drosophila* S2 cells. These experiments were performed in triplicate, all on the same day, in the Wittkopp lab. For each reporter gene, a t-test was used to compare expression of the reporter gene with and without *miR-8* overexpression, and expression levels were normalized to expression of the reporter gene in the absence of *miR-8* overexpression. Data from these experiments in S2 cells are shown in **Supplementary Figure S5**. Sequences of 3’ UTRs used in these experiments, including annotations of putative *miR-8* binding sites and information about their conservation, are provided in **Supplementary File S1**.

## Results and Discussion

### Gain-of-function screen: 48 of 166 miRNAs are able to alter body pigmentation in adult flies

To identify miRNAs that can affect development of body pigmentation in adult *D. melanogaster*, we used the UAS/Gal4 system (Brand and Perrimon 1993) to overexpress 166 miRNAs in developing flies and assessed their effects on pigmentation (**Supplementary Table S1**). [Crosses designed to overexpress 4 additional miRNAs were also set up, but failed to produce progeny (**Supplementary Table S1, gray highlight**).] Of the 166 miRNAs tested, 86% were confidently annotated as miRNAs in *D. melanogaster* by Kozomara et al. (2019) and 98% were included in the collection of UAS-miRNA lines described in Schertel et al. (2012). We used *pannier*-Gal4 (*pnr*-Gal4) to overexpress each miRNA. We chose this driver because it expresses Gal4 in dorsal epidermal cells at the pupal stages during which adult pigmentation develops (Wittkopp et al. 2002; Rogers et al. 2014; Kalay et al. 2016).

For 19 of the 166 miRNAs tested, fewer adult progeny inherited both the Gal4 driver and UAS construct than expected by chance assuming Mendelian inheritance (one-sided binomial test, p < 0.01, **Supplementary Table S1, column K**), suggesting that overexpression of those miRNAs reduced viability. 13 of these 19 miRNAs were excluded from our analysis because the crosses produced 2 or fewer adult progeny with both the UAS and Gal4 transgenes, suggesting that overexpression of these miRNAs was lethal (or nearly so) (**Supplementary Table S1, red font**).

For each of the remaining 153 miRNAs, pigmentation was scored in 2 to 45 males (average = 9.41) and 1 to 35 females (average = 10.08) carrying both the UAS and Gal4 constructs (**Supplementary Table S1, columns L and M**). This scoring was performed separately for the thorax, anterior abdomen (abdominal segments A2-A4), and posterior abdomen (abdominal segments A5-A6) by visually comparing pigmentation of the dorsal cuticle in the *pnr-Gal4* expression domain to pigmentation in the flanking (more lateral) regions of the same animal, which lack Gal4 expression (**Figure 1B**). For each region, in each sex, if pigmentation in the *pnr-Gal4* expression domain was lighter or darker than the flanking cuticle, the fly was scored as being “lightened” or “darkened” by the miRNA, respectively (**Figure 1C**). The penetrance of these phenotypes was calculated for each miRNA in each sex as the number of flies showing altered pigmentation divided by the total number of flies scored for that genotype. Flies with unaltered pigmentation in the *pnr-Gal4* expression domain were considered “unaffected” (**Figure 1C**).

We found that overexpressing 105 of these 153 miRNAs did not cause a visible change in pigmentation in the *pnr-Gal4* expression domain with a penetrance ≥ 50% in any body region in either sex, suggesting they are not able to alter pigmentation development. By contrast, overexpressing each of the 48 remaining miRNAs caused a visible change in pigmentation in the *pnr-Gal4* expression domain in at least one body region of at least one sex with a penetrance ≥ 50%. As shown in **Table 1**, 25 of these 48 miRNAs induced lighter pigmentation in one or more body regions, 29 induced darker pigmentation in one or more body regions, and 6 caused lighter pigmentation in some body regions but darker pigmentation in others.

**Table 1.**
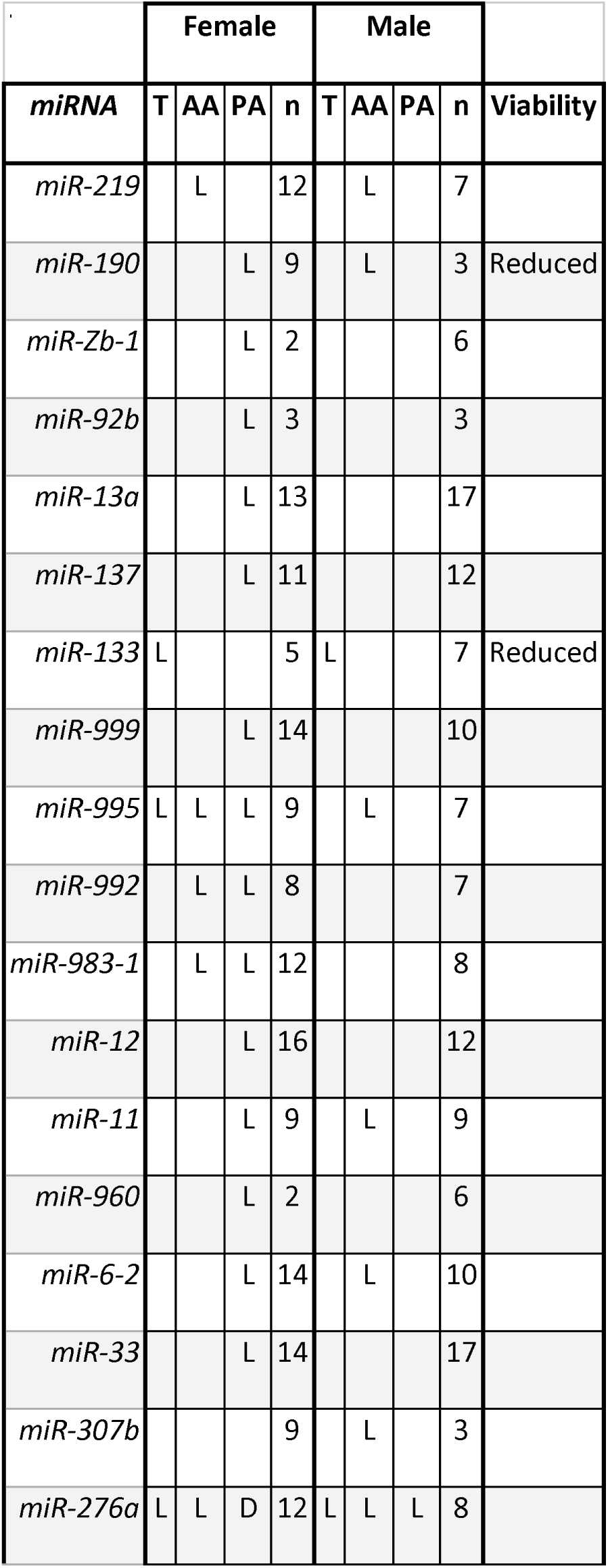
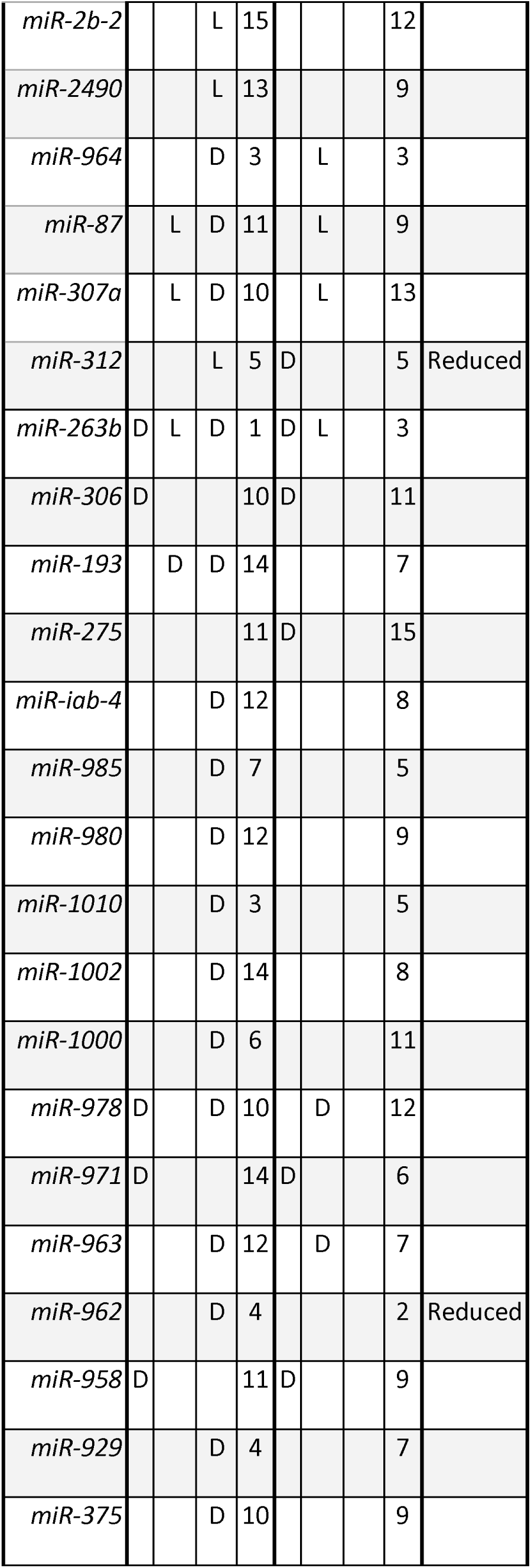
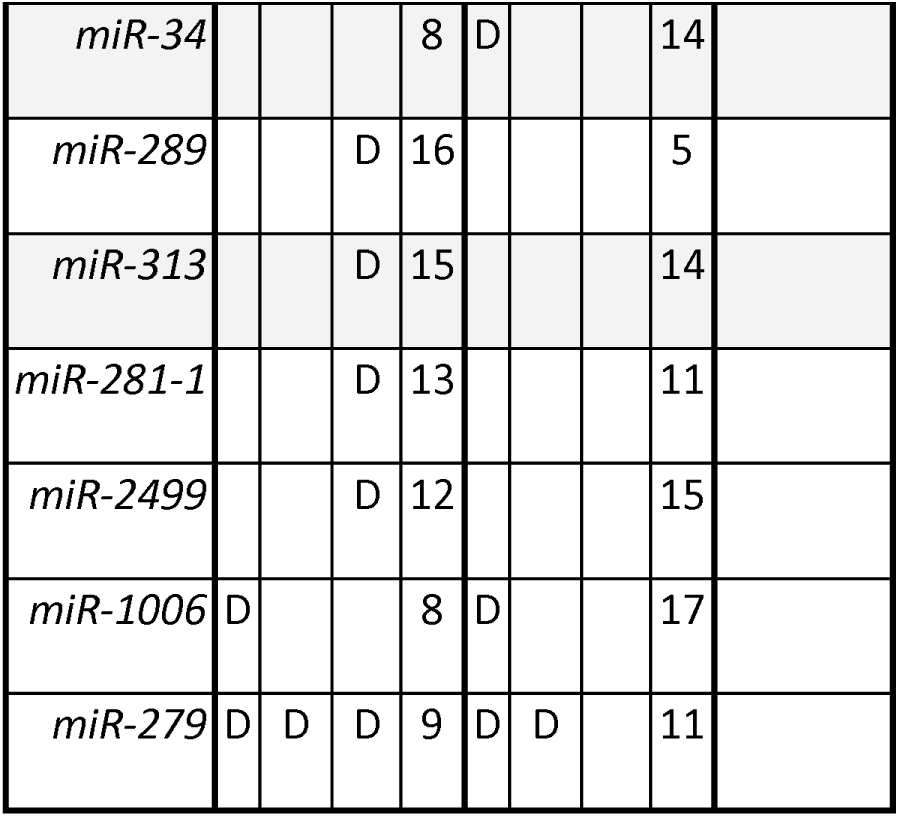
Effects of miRNA overexpression on adult body pigmentation. Data are shown for the 48 of 166 miRNAs tested that caused a visible change in pigmentation within the *pnr-Gal4* expression domain of the thorax (T), anterior abdomen (AA), or posterior abdomen (PA) at a penetrance of at least 50% in females or males. “L” indicates that pigmentation was lightened and “D” indicates that it was darkened. The number of flies scored (n) for each sex is also shown. Cases where overexpression of the miRNA with the *pnr-Gal4* driver resulted in significantly reduced viability are also shown. For miRNAs without effects on pigmentation, the number of individuals scored and information about viability are shown in **Supplementary Table S1**.

The female posterior abdomen was the most often affected body region, with 39 of these 48 miRNAs altering its pigmentation (**Table 1**, **Figure 2A**). In 23 of these 39 cases (59%), the posterior abdomen was the only body region with pigmentation affected by overexpressing a miRNA (**Table 1**). These sex-specific effects in the posterior abdomen are consistent with pigmentation of abdominal segments A5 and A6 in *D. melanogaster* being sexually dimorphic. Effects in the female posterior abdomen were seen much more often in segment A6 than in segment A5 (**Figure 2B; Supplementary Table S2**). Interestingly, for pigmentation, prior work has also shown A6 in females to be the abdominal segment most sensitive to genetic variation (Kopp et al. 2003; Salomone et al. 2013; Dembeck, Huang, Magwire, et al. 2015; Yassin et al. 2016), changes in transcription factor activity (Rogers et al. 2014; Kalay et al. 2016), loss of *miR-8* or *miR-33* expression (Kennell et al. 2012; Clerbaux et al. 2021), and temperature (Gibert et al. 2004; Gibert et al. 2007). In males, only one of the miRNAs tested (*miR-276a*) visibly altered (lightened) pigmentation in the posterior abdomen, although we note that miRNAs darkening pigmentation would be hard to detect because segments A5 and A6 are already heavily melanized in wild-type males.

In the thorax, 10 miRNAs altered pigmentation in females and 11 altered pigmentation in males, with 8 having effects in both sexes, all of which were in the same direction (2 lightening and 6 darkening) (**Table 1**, **Figure 2**). This difference in results between males and females is unexpected because thorax pigmentation is not considered to be sexually dimorphic in *D. melanogaster*. It is possible that the 5 miRNAs with sex-specific overexpression phenotypes in the thorax might have effects in the other sex that were missed due to the subtlety of the phenotype, small sample sizes, or variable penetrance.

In the anterior abdomen (A2-A4), 10 miRNAs were observed to alter pigmentation in females and 14 were observed to alter pigmentation in males, with 7 having effects in both sexes, all of which were in the same direction (6 lightning and 1 darkening) (**Table 1**, **Figure 2**). As with the thorax, pigmentation in abdominal segments A2-A4 is not generally considered sexually dimorphic (although sex-specific differences in plasticity have been observed (Gibert et al. 2009)), suggesting that the miRNAs with sex-specific effects seen here (3 female-only and 7 male-only) might also have effects in the opposite sex that we did not observe.

As noted above, 42 of the 48 (87.5%) miRNAs with overexpression phenotypes either had the same effect (lightening or darkening) in both sexes or only affected pigmentation in one sex (**Table 1**). For the 6 miRNAs where this was not the case, 5 showed effects on the female posterior abdomen that were in the opposite direction from the effects observed elsewhere in females or in the males (**Table 1**). In the sixth case, overexpressing *miR-276a* was observed to lighten pigmentation in all regions in both sexes except for the female posterior abdomen, where it was observed to broaden the dark pigmentation. These differences in the effects of miRNA overexpression among body regions and sexes could result from differences in the presence of transcripts from miRNA target genes in different regions and sexes that are available to be repressed by the miRNA.

Of the 3 miRNAs previously shown to affect pigmentation development in *D. melanogaster*, we observed overexpression phenotypes consistent with prior work for two of them: *miR-193* (Tian et al. 2024) and *miR-33* (Clerbaux et al. 2021) (**Table 1**). Interestingly, we did not observe a visible effect on pigmentation when overexpressing *miR-8*, which was the first miRNA shown to affect pigmentation in *Drosophila melanogaster* (Kennell et al. 2012). One possible explanation for this finding is that *miR-8* might already be expressed at high levels in the female A6 segment, where loss of *miR-8* has been shown to alter pigmentation.

Observation notes from each of the 6387 flies scored in this overexpression screen, plus information about 580 control flies (87 *yellow* mutants, 241 *ebony* mutants, and 252 *UAS-ebony-RNAi/pnr-Gal4* flies) and 98 flies expressing a competitive inhibitor for *mir-8* are provided in **Supplementary Table S2.** Developmental defects affecting traits other than pigmentation that were observed in this screen are also described in this table.

### Loss-of-function screen: 22 out of 38 miRNA inhibitors altered body pigmentation in adult flies

The overexpression screen described above identified miRNAs that are sufficient to affect pigmentation, but these miRNAs may or may not be expressed in the right time or place to affect normal adult pigmentation development. To identify miRNAs that normally play a role in pigmentation development, we used competitive inhibition to reduce the activity of 41 miRNAs in the same *pnr-Gal4* expression domain used for the overexpression screen. These 41 miRNAs included 37 of the 48 miRNAs found to have effects on pigmentation with ≥50% penetrance in our overexpression screen as well as *miR-8*, *miR-965, miR-6-1,* and *miR-6-3*. We included *miR-8* (whose overexpression darkened pigmentation in the female posterior abdomen with a penetrance 37%) as a positive control because prior work showed that *miR-8* loss-of-function affects pigmentation development (Kennell et al. 2012). *miR-965*, which was previously shown to affect abdominal development (Verma and Cohen 2015) was included because of the distinctiveness of its overexpression phenotype even though it was only 31% penetrant. *miR-6-1* and *miR-6-3* (whose overexpression did not alter pigmentation) were included because they are affected by the competitive inhibitor used for *miR-6-2* (which altered pigmentation in the overexpression screen) (**Table 1**, **Figure 2B**), as described below.

To reduce activity of these 41 miRNAs, we used 38 previously constructed miRNA “sponge” lines, which contain UAS sequences controlling expression of a reporter gene containing a 3’ UTRs with 20 recognition sites for a specific miRNA(s) (Fulga et al. 2015) (**Figure 1D**). 36 of these miRNA sponge lines targeted a single miRNA, whereas the other 2 each targeted multiple miRNAs (*miR-6-1/miR-6-2/miR-6-3* and *miR-2b-1/mir-2b-2*) (**Supplementary Table S1**). These miRNA sponge transcripts compete with native mRNA targets for binding to a specific miRNA(s), effectively reducing activity of the targeted miRNA(s) in the *pnr-Gal4* expression domain during pigmentation development.

We suspected that competitive inhibition of native miRNAs might cause more subtle effects on pigmentation than overexpression, so we set up crosses to test the effects of each sponge on pigmentation at two different temperatures that are known to promote darker (18°C) or lighter (28°C) body pigmentation (Gibert et al. 2004). We reasoned that the effects of miRNAs lightening pigmentation might be easier to detect on more darkly pigmented flies, whereas the effects of miRNAs darkening pigmentation might be easier to detect on more lightly pigmented flies. We again scored pigmentation in the thorax, anterior abdomen, and posterior abdomen of males and females as lightened, darkened, or unaffected (**Figure 1D**), both by comparing pigmentation within and outside the *pnr-Gal4* expression domain in the dorsal cuticle as well as comparing each inhibition phenotype to a genetically matched control line expressing a “scrambled” sponge that does not inhibit any miRNA in the *D. melanogaster* genome (Fulga et al. 2015). This secondary control proved to be important because some of the UAS-RNAi-sponge lines showed evidence of low-level sponge expression outside the *pnr-Gal4* expression domain.

For the flies reared at 18°C, we scored 4 to 41 males (average 11.4) and 6 to 49 females (average 12.2), for all miRNAs except *miR-971*, as the cross for that miRNA failed to produce any progeny (**Supplementary Table S1**). For the flies reared at 28°C, we scored 2 to 35 males and 1 to 38 females, with an average of 11.2 males and 10.6 females, for all miRNAs (**Supplementary Table S1**). Inhibiting activity of these miRNAs did not significantly reduce viability in any case (one-sided binomial test, all p > 0.01, **Supplementary Table S1, columns AD and AU**). Observation notes from each of the 3499 flies scored with miRNA activity inhibited, plus information from 246 control flies expressing the “scrambled” version of a miRNA sponge, are provided in **Supplementary Table S3.** Observations of developmental defects affecting traits other than pigmentation are also included in this table.

We found that 22 of the 38 miRNA sponges caused visible changes in pigmentation with a penetrance of ≥50% in at least one body region of at least one sex at one or both rearing temperatures. (**Table 2**, **Supplementary Table S1).** As with the overexpression screen, reducing activity of a miRNA affected pigmentation most often in the female posterior abdomen (n = 20) (**Table 2**). 15 miRNAs lightened pigmentation in one or more body regions in one or both sexes, whereas 6 darkened it (**Table 2**). The remaining 2 miRNAs lightened pigmentation in females in one region, but darkened it in another (**Table 2**). miRNA sponges for *miR-8*, *miR-279*, *miR-12*, and *miR-33* produced visible pigmentation phenotypes at >50% penetrance when reared in both temperatures, whereas the rest of the miRNA sponges caused pigmentation phenotypes seen only in flies reared at only one of the temperatures, most often at 28°C (**Table 2**).

**Table 2.**
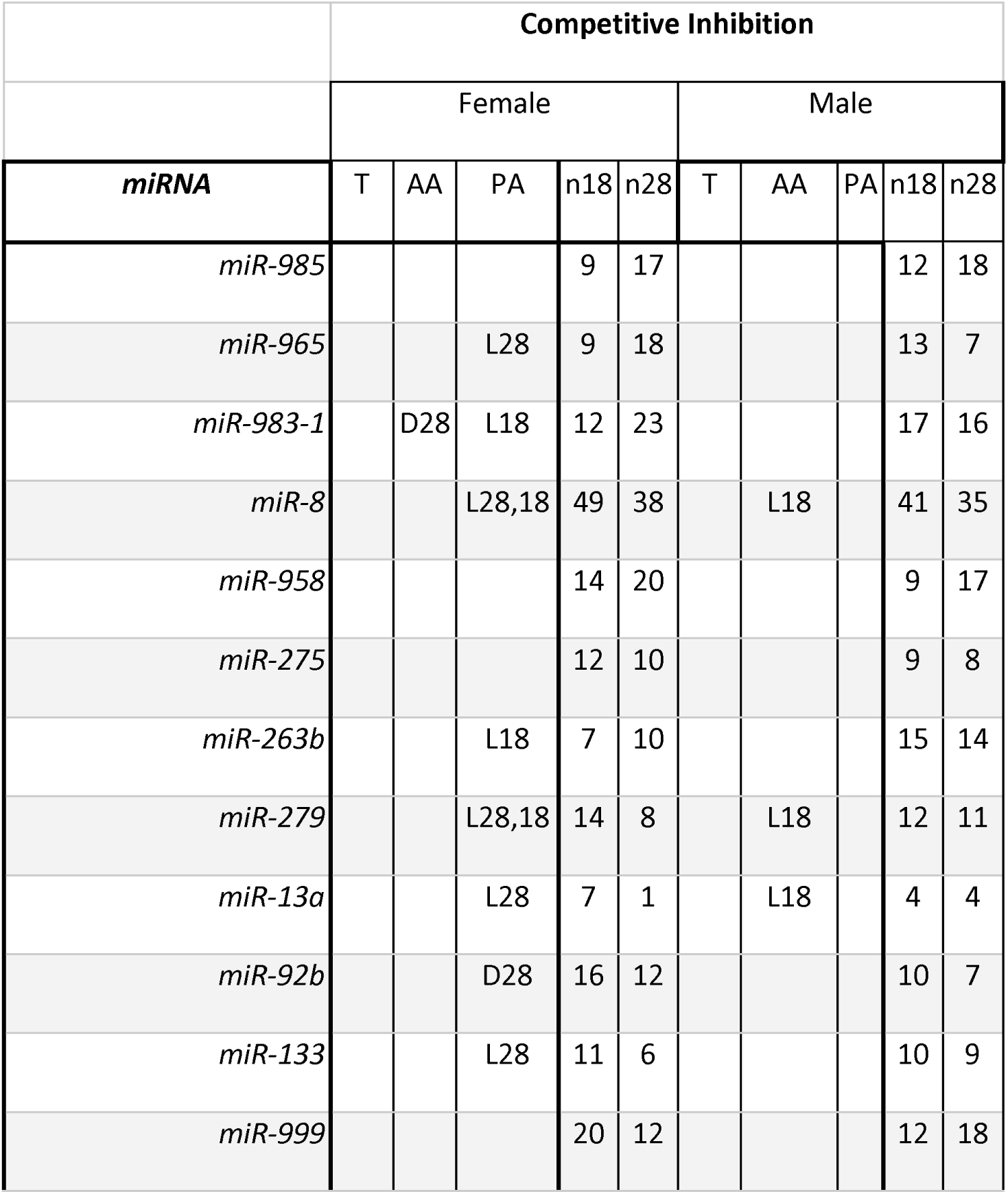
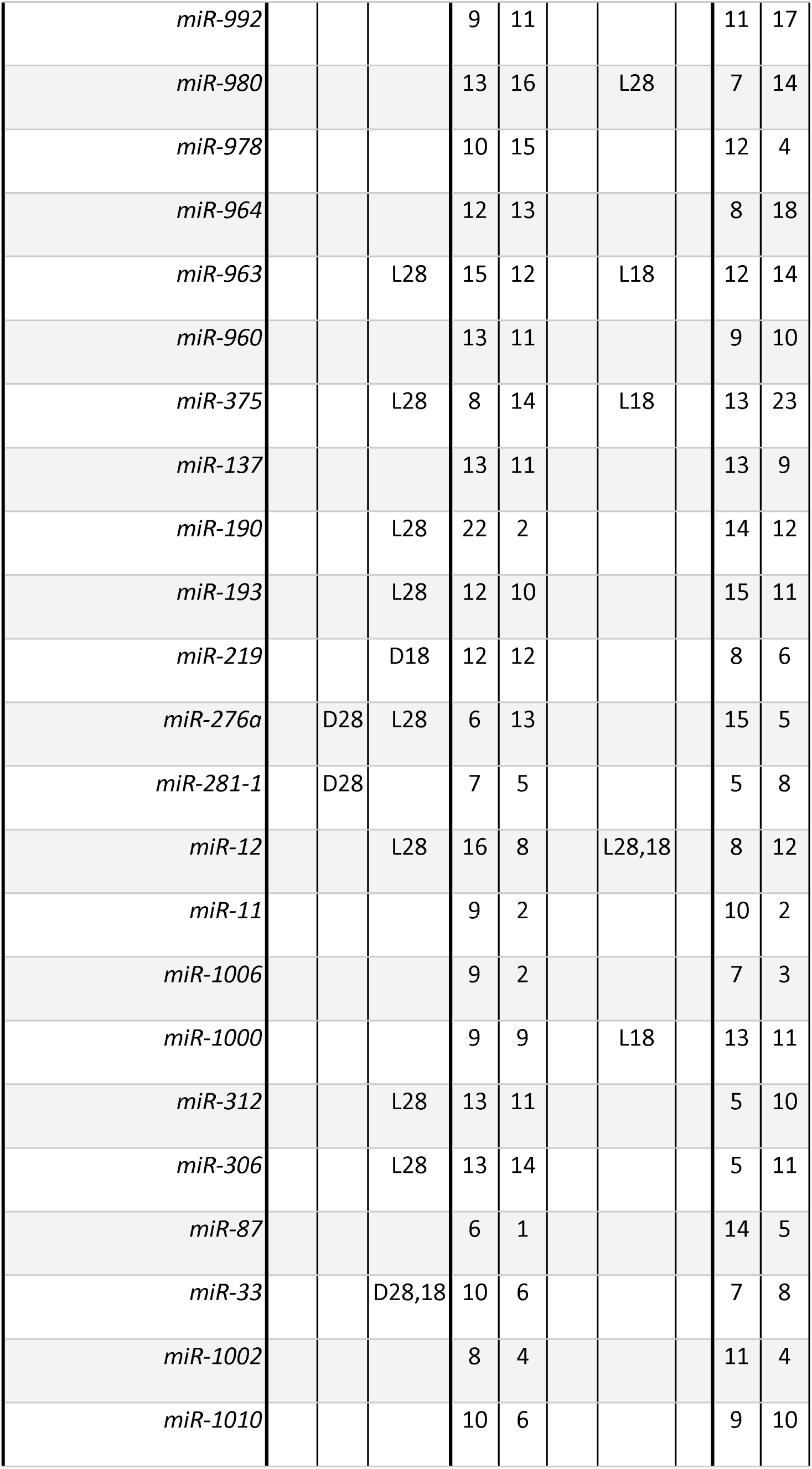
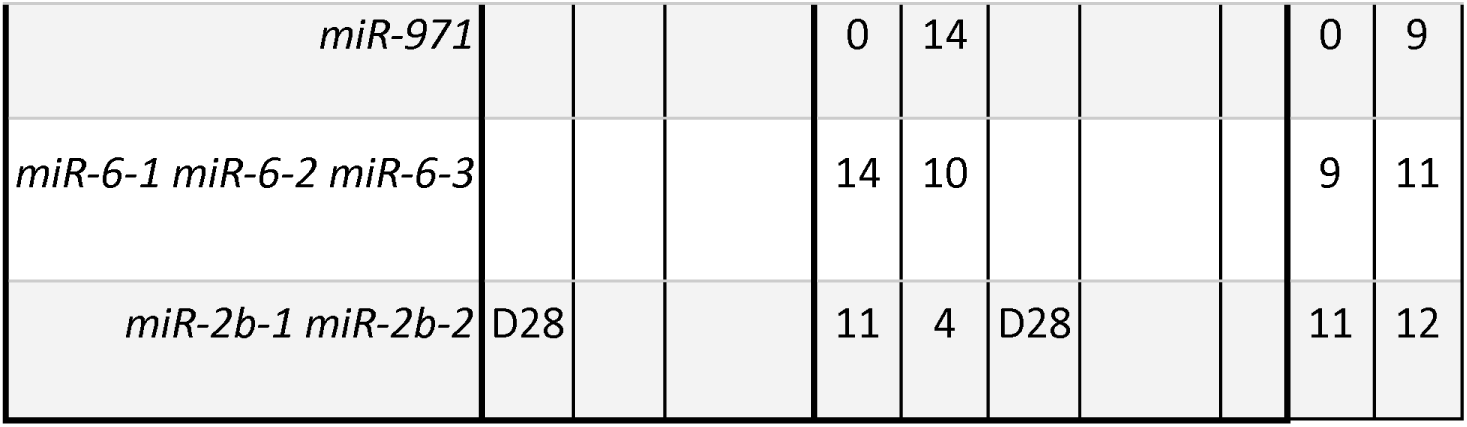
Effects of reducing miRNA activity on adult body pigmentation. Data are shown for the 41 miRNAs competitively inhibited by 38 miRNA sponge constructs. *miR-6-1*, *miR-6-2*, and *miR-6-3* have the same seed sequence, so were inhibited by a single sponge construct. Similarly, miR-2b-2 and miR-2b-2 also share a seed sequence and were both inhibited by a single sponge construct. Pigmentation was scored in the thorax (T), anterior abdomen (AA), and posterior abdomen (PA) in females or males. “L” indicates that pigmentation was lightened, and “D” indicates that it was darkened, at a penetrance of at least 50% in the *pnr-Gal4* expression domain. The subscript(s) (18 or 28) indicate the rearing temperature at which the effect was seen. The number of females and males scored after rearing at 18°C (n_18_) and 28°C (n_28_) are also shown. For *miR-971*, phenotypes were only examined for flies reared at 28°C because the 18°C cross failed to produce any offspring. A marginally significant reduction in viability was also observed for flies with reduced *miR-279* at 28°C (binomial exact test, p = 0.046, **Supplementary Table S1**).

| Table 2 |  | Competitive Inhibition |  |  |  |  |  |  |  |  |  |
| --- | --- | --- | --- | --- | --- | --- | --- | --- | --- | --- | --- |
|  |  | Female |  |  |  |  | Male |  |  |  |  |
| *miRNA* |  | T | AA | PA | n18 | n28 | T | AA | PA | n18 | n28 |
| *miR-985* |  |  |  |  | 9 | 17 |  |  |  | 12 | 18 |
| *miR-965* |  |  |  | L28 | 9 | 18 |  |  |  | 13 | 7 |
| *miR-983-1* |  |  | D28 | L18 | 12 | 23 |  |  |  | 17 | 16 |
| *miR-8* |  |  |  | L28,18 | 49 | 38 |  | L18 |  | 41 | 35 |
| *miR-958* |  |  |  |  | 14 | 20 |  |  |  | 9 | 17 |
| *miR-275* |  |  |  |  | 12 | 10 |  |  |  | 9 | 8 |
| *miR-263b* |  |  |  | L18 | 7 | 10 |  |  |  | 15 | 14 |
| *miR-279* |  |  |  | L28,18 | 14 | 8 |  | L18 |  | 12 | 11 |
| *miR-13a* |  |  |  | L28 | 7 | 1 |  | L18 |  | 4 | 4 |
| *miR-92b* |  |  |  | D28 | 16 | 12 |  |  |  | 10 | 7 |
| *miR-133* |  |  |  | L28 | 11 | 6 |  |  |  | 10 | 9 |
| *miR-999* |  |  |  |  | 20 | 12 |  |  |  | 12 | 18 |

|  |  |  |  |  |  |  |  |  |  |  |
| --- | --- | --- | --- | --- | --- | --- | --- | --- | --- | --- |
| *miR-992* |  |  |  | 9 | 11 |  |  |  | 11 | 17 |
| *miR-980* |  |  |  | 13 | 16 |  | L28 |  | 7 | 14 |
| *miR-978* |  |  |  | 10 | 15 |  |  |  | 12 | 4 |
| *miR-964* |  |  |  | 12 | 13 |  |  |  | 8 | 18 |
| *miR-963* |  |  | L28 | 15 | 12 |  | L18 |  | 12 | 14 |
| *miR-960* |  |  |  | 13 | 11 |  |  |  | 9 | 10 |
| *miR-375* |  |  | L28 | 8 | 14 |  | L18 |  | 13 | 23 |
| *miR-137* |  |  |  | 13 | 11 |  |  |  | 13 | 9 |
| *miR-190* |  |  | L28 | 22 | 2 |  |  |  | 14 | 12 |
| *miR-193* |  |  | L28 | 12 | 10 |  |  |  | 15 | 11 |
| *miR-219* |  |  | D18 | 12 | 12 |  |  |  | 8 | 6 |
| *miR-276a* |  | D28 | L28 | 6 | 13 |  |  |  | 15 | 5 |
| *miR-281-1* |  | D28 |  | 7 | 5 |  |  |  | 5 | 8 |
| *miR-12* |  |  | L28 | 16 | 8 |  | L28,18 |  | 8 | 12 |
| *miR-11* |  |  |  | 9 | 2 |  |  |  | 10 | 2 |
| *miR-1006* |  |  |  | 9 | 2 |  |  |  | 7 | 3 |
| *miR-1000* |  |  |  | 9 | 9 |  | L18 |  | 13 | 11 |
| *miR-312* |  |  | L28 | 13 | 11 |  |  |  | 5 | 10 |
| *miR-306* |  |  | L28 | 13 | 14 |  |  |  | 5 | 11 |
| *miR-87* |  |  |  | 6 | 1 |  |  |  | 14 | 5 |
| *miR-33* |  |  | D28,18 | 10 | 6 |  |  |  | 7 | 8 |
| *miR-1002* |  |  |  | 8 | 4 |  |  |  | 11 | 4 |
| *miR-1010* |  |  |  | 10 | 6 |  |  |  | 9 | 10 |
| *miR-971* |  |  |  | 0 | 14 |  |  |  | 0 | 9 |
| *miR-6-1 miR-6-2 miR-6-3* |  |  |  | 14 | 10 |  |  |  | 9 | 11 |
| *miR-2b-1 miR-2b-2* | D28 |  |  | 11 | 4 | D28 |  |  | 11 | 12 |

### 15 miRNAs affecting pigmentation development in adult D. melanogaster

As described above, inhibiting the activity of a miRNA tests whether its native expression is necessary for pigmentation development, whereas overexpressing a miRNA tests whether it is sufficient to alter pigmentation. Therefore, miRNAs that displayed opposite effects in our two screens, particularly with effects in the same body region for both screens, represent our highest confidence candidates for miRNAs affecting pigmentation development. To identify such cases, we compared the results from our gain-of-function and loss-of-function screens. We focus our discussion on the effects of overexpressing and inhibiting miRNAs in females using the inhibition data from 28°C (**Figure 3**), where the largest number of effects were seen (**Table 1**, **Table 2**). However, in the Supplementary Materials, we show the same comparisons for males using inhibition phenotypes seen at 28°C (**Supplementary Figure S1**) as well as for females (**Supplementary Figure S2**) and males (**Supplementary Figure S3**) using inhibition phenotypes seen at 18°C.

**Figure 3.**
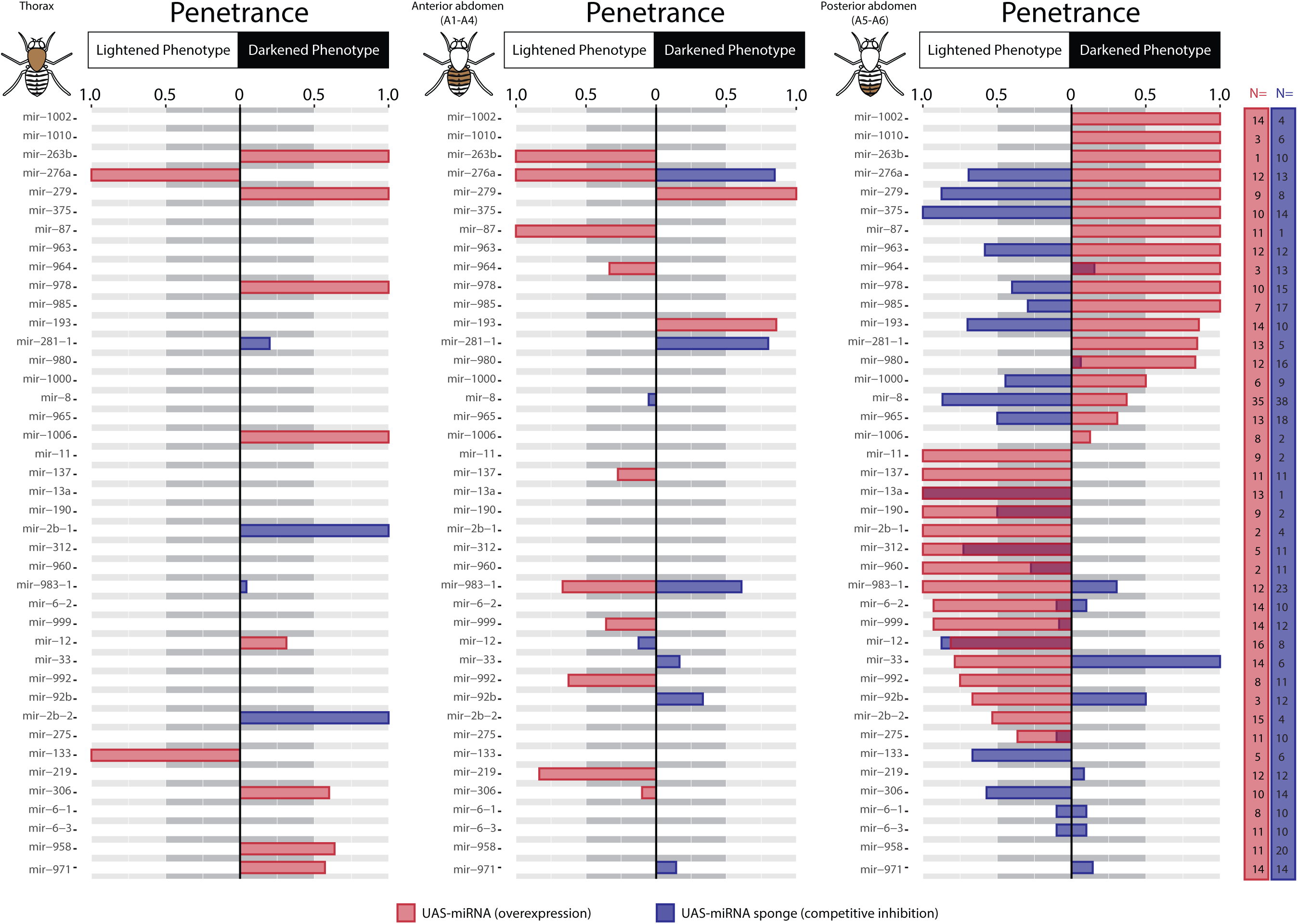
Pigmentation phenotypes caused by overexpression and competitive inhibition of miRNAs. From left to right, the plots show the penetrance of “lightened” or “darkened” pigmentation phenotypes observed in the thorax, anterior abdomen, and posterior abdomen of female flies. Penetrance is defined by the number of flies classified as having that phenotype divided by the number of flies scored with the same genotype. Consequently, penetrance ranges from 0 (no flies observed displayed the phenotype) to 1 (all flies observed display the phenotype). Penetrance of lightened phenotypes is plotted extending to the left from the “0” axis, while penetrance of darkened phenotypes is plotted extending to the right. Phenotypes resulting from overexpression are shown by red bars, whereas phenotypes resulting from competitive inhibition (at 28°C) are shown by blue bars. The number of flies scored for each genotype with altered miRNA expression is shown on the right of the figure under “N=” using the same color coding as in the penetrance plots (i.e., red = overexpression; blue = competitive inhibition). Note that *miR-6-1*, *miR-6-2*, and *miR-6-3* share a seed sequence and thus were simultaneously inhibited by a single miRNA sponge, resulting in the same competitive inhibition data for these 3 miRNAs. *miR-2b-1* and *miR-2b-2* also share a seed sequence and were inhibited by a single miRNA sponge. A version of this figure using data from males reared at 28°C is provided as **Supplementary Figure S1**. Versions of this figure for females and males using data from the competitive inhibition crosses maintained at 18°C are provided as **Supplementary Figure S2** and **Supplementary Figure S3**, respectively. Summary data from all screen crosses is available in **Supplementary Table S1**.

In the thorax, none of the miRNAs tested showed opposite effects on pigmentation in the gain-of-function and loss-of-function screens. However, inhibiting *miR-2b-1* and *miR-2b-2* together caused fully (100%) penetrant darker pigmentation phenotypes in the thorax, whereas overexpressing either of these miRNAs lightened pigmentation in the female posterior abdomen with penetrance ≥50% (**Figure 3**). This pattern of effects in different tissues could be explained by endogenous expression of *miR-2b-1* and/or *miR-2b-2* being higher in the thorax than in the female posterior abdomen. Overexpressing *miR-276a* or *miR-279* also caused fully penetrant pigmentation phenotypes in the thorax with opposite changes in pigmentation (with ≥50% penetrance) caused by inhibition in one (*miR-279*) or both (*miR-276a*) regions of the female abdomen examined (**Figure 3**). Manipulating activity of *miR-306* also showed opposite effects on pigmentation in the thorax and in the female posterior abdomen, but with penetrance closer to 50% (**Figure 3**).

In the abdomen, overexpressing or inhibiting 2 miRNAs (*miR-276a* and *miR-983-1*) had opposite effects on pigmentation with penetrance ≥50% in the anterior segments (A2-A4), suggesting they play a role in normal pigmentation development (**Figure 3**). In the posterior abdomen (segments A5 and/or A6), 7 miRNAs (miR−276a, miR-279, miR-375, miR-963, miR-193, miR-33, and *miR-92b*) showed opposite phenotypes with a penetrance ≥50% in both the overexpression and inhibition screens (**Figure 3**), including 2 of the 3 miRNAs previously shown to affect adult pigmentation in *D. melanogaster*: *miR-193* (Tian et al. 2024) and *miR-33* (Clerbaux et al. 2021). The third previously identified miRNA, *miR-8* (Kennell et al. 2012), also showed opposing phenotypes in the gain-of-function and loss-of-function screens, but the penetrance of the overexpression phenotype in the female posterior abdomen was only 37% (**Figure 3; Supplementary Table S1**). In all three cases, the effects described here are consistent with previously published results. *miR-965, miR-1000,* and *miR-978* showed opposite pigmentation phenotypes in the female posterior abdomen in the two screens with the inhibition phenotype having at least 40% penetrance, suggesting they are also worthy of further study.

Taken together, this analysis identified 8 miRNAs with the strongest evidence of affecting normal pigmentation development in adult *D. melanogaster* (i.e., complementary effects of overexpression and inhibition in the same body region with penetrance ≥50%.): *miR-276a, miR- 983-1, miR-279, miR-375, miR-963, miR-193, miR-33,* and *miR-92b*. We also consider *miR-8*, *miR-2b-1, miR-2b-2, miR-306, miR-965, miR-1000,* and *miR-978* to be strong candidates as miRNAs affecting pigmentation for the reasons discussed above.

Overexpression and inhibition phenotypes for 4 of these 15 miRNAs (*miR-33, miR-279, miR-92b,* and *miR-276a*) are shown in **Figure 4**, and the overexpression phenotypes for *miR-193* and *miR-978* are shown in **Figure 2B**. The *miR-276a* phenotypes are particularly striking (**Figure 4G**). Overexpressing *miR-276a* lightened pigmentation in females within the *pnr-Gal4* expression domain in the anterior abdomen (A2-A4) as well as the more anterior segment in the posterior region (A5), but darkened pigmentation in the most posterior segment, A6 (**Figure 4G,4G’, and 4G’’**). Reducing activity of *miR-276a* through competitive inhibition had opposing effects in females, with pigmentation in the *pnr-Gal4* expression domain getting darker in segments A2-A5 and lighter in segment A6 (**Figure 4H**). In males, overexpressing *miR-276a* caused a visible lightening in all pigmented abdominal segments (A2-A6) (**Figure 4G’’’**).

**Figure 4.**
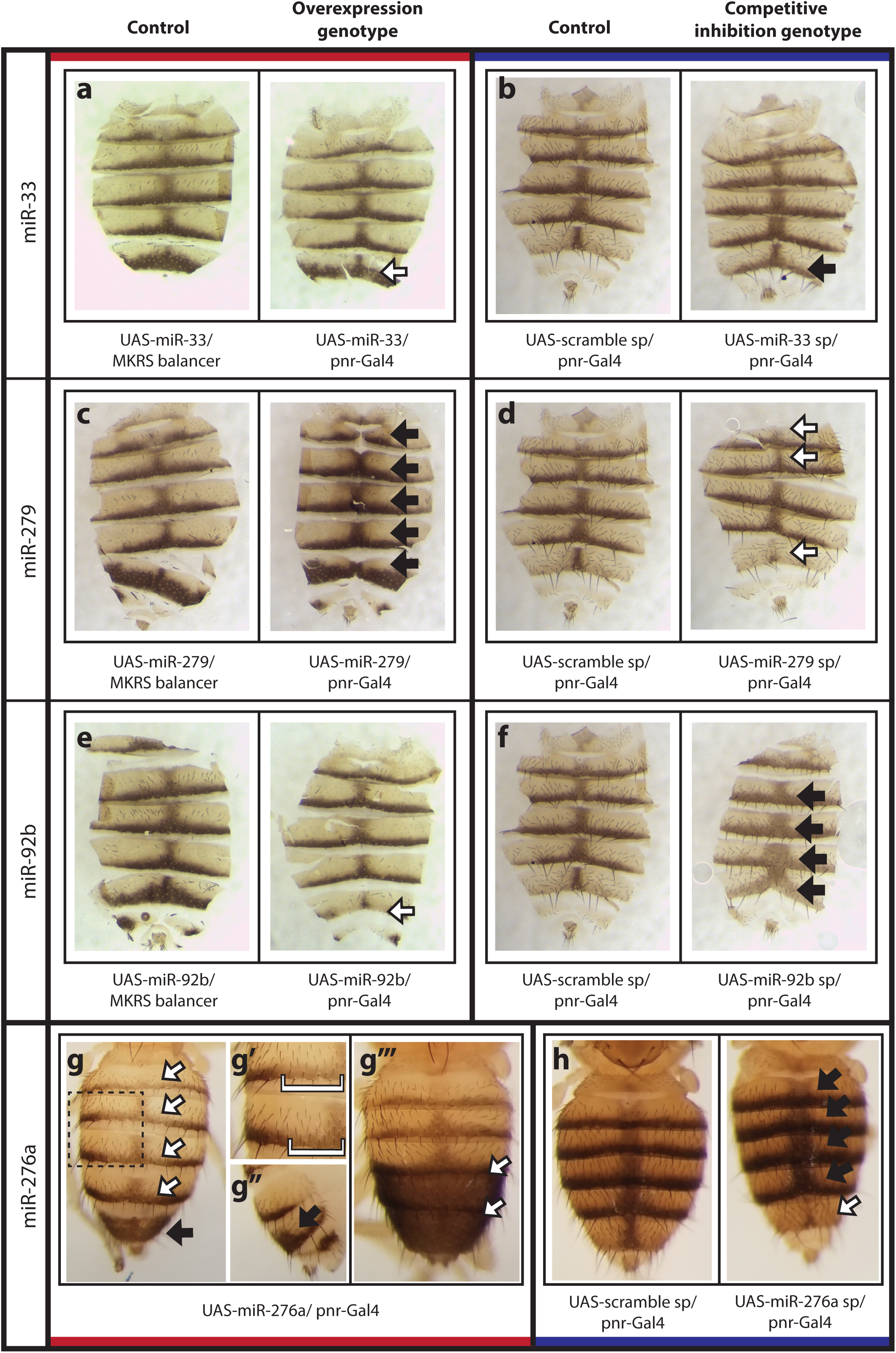
Examples of miRNAs that are both sufficient to alter abdominal pigmentation and necessary for normal pigmentation development. All images show dorsal abdominal cuticle from 3-5 day old adult flies. White arrows and brackets indicate lightened pigmentation in the *pnr-Gal4* expression domain. Black arrows represent areas of darkened pigmentation in the *pnr-Gal4* expression domain. (**A-F)** Dissected and mounted dorsal abdominal cuticles of female flies. For each panel, flies that either overexpress (**A, C, E**) or competitively inhibit (**B, D, F**) the target miRNA are on the right, with matched control flies on the left. Data is shown for *miR-33* (**A, B**), *miR-279* (**C, D**), and *miR-92b* (**E, F**). (**G,H)** Whole abdomens of flies that carry transgenes to overexpress (**G-G’’’**) or competitively inhibit (**H**) *miR-276a*. (**G)** Dorsal view of female abdomen overexpressing *miR-276a* in the *pnr-Gal4* expression domain. (**G’)** Enlarged view of region indicated by the dashed box in panel G. Areas of lightened pigmentation are marked by brackets. (**G’’)** Right lateral view of abdominal segments A5 and A6 in the same fly shown in panels **G** and **G’**. Black arrow points to the sharp increase in melanization outside of the *pnr-Gal4* expression domain. (**G’’’)** Dorsal view of a male fly overexpressing *miR-276a*. White arrows point to reduced melanization of segments A5 and A6. (**H)** Whole abdomen from female fly expressing the scrambled control miRNA sponge (left). Whole abdomen from female fly expressing the miRNA sponge targeting *miR-276a* (right). White arrow shows lightened pigmentation in segment A6; black arrows show darkened pigmentation in segments A2-A5. The genotype of each fly is shown beneath each panel. The red bars at the top and bottom of the figure indicate flies from the overexpression screen, and the blue bars at the top and bottom of the figure indicate flies from the competitive inhibition screen.

As described above, deletion of *miR-8* <u>(Kennell et al. 2012)</u> and *miR-33* <u>(Clerbaux et al.</u> <u>2021)</u> have previously been shown to affect pigmentation in *D. melanogaster,* and deletion of *miR-193* has been shown to affect pigmentation in butterflies <u>(Tian et al. 2024)</u>. We also examined pigmentation in flies homozygous for deletions of *miR-92b*, *miR-193*, and *miR-276* using lines produced by <u>Chen et al. (2014)</u> and obtained from the Bloomington Drosophila Stock Center. We found that deleting any of these three miRNAs was sufficient to alter body pigmentation, with the deletion of *miR-92b* lightening pigmentation and the deletion of *miR-193* or *miR-276* darkening pigmentation. Phenotypes for these deletion mutants are shown and compared to the phenotypes resulting from expression of miRNA sponge constructs in **Supplementary Figure S4**. Both similarities and differences were observed, and the differences might result from the competitive inhibition sponges reducing miRNA activity only in the *pnr-Gal4 expression domain*, which is a subset of cells during a subset of development, and the flies homozygous for the knockout allele missing the microRNA in all tissues at all developmental stages. These differences might also be caused by off-target effects of the sponge or other unknown effects.

### Identifying potential target genes for 15 miRNAs affecting pigmentation

miRNAs repress expression of their target genes by binding to short 6-8 base sequences in the 3’ UTRs that are complementary to a region in the miRNA known as the “seed” (Bartel 2018) (**Figure 1A**). This seed region of a miRNA can be identified by its position in the primary miRNA transcript, allowing potential targets of miRNAs to be identified computationally by scanning the genome for seed matches in the 3’ UTRs of protein-coding genes (Enright et al. 2003; Lewis et al. 2003). However, the presence of a sequence complementary to the seed is only one of the factors that allows a miRNA to pair with its target, and these computational methods are known to produce false positives and false negatives (Pinzón et al. 2017). Other factors that can influence miRNA binding include (a) that the position of the seed sequence and/or structure of the 3’UTR can make the seed site inaccessible (Agarwal et al. 2018), (b) that even imperfect seed matches and pairing with bases outside the seed region of the mature miRNA can lead to repression of target genes (Broughton et al. 2016), and (c) that the miRNA, target gene, and all components of the RISC complex might not be expressed in the same cells at the same time. Nonetheless, knowing the location of 3’ UTR sequences complementary to an miRNAs seed sequence remains a helpful tool for generating hypotheses about possible targets of miRNAs.

To identify possible miRNA-target gene interactions likely to affect pigmentation, we searched for potential interactions between the seed sequences of the 15 miRNAs implicated in pigmentation development and the 3’UTR sequences of 93 genes shown previously to alter adult body pigmentation in *D. melanogaster* (**Supplementary Table S4**). These 93 genes were identified by first using gene ontology annotations in Flybase (Thurmond et al. 2019) and then manually filtering out genes lacking experimental evidence that they affect adult body pigmentation (see Materials and Methods). Pigmentation genes identified in large-scale screens or GWAS studies of pigmentation in *D. melanogaster* (Rogers et al. 2014; Dembeck, Huang, Carbone, et al. 2015; Kalay et al. 2016) were added to the list only if additional experimental evidence that the gene affects pigmentation existed.

For each of the 93 genes, we described its role in pigmentation development as “darkens” if the gene had been experimentally shown to be either necessary or sufficient for the development of dark pigmentation, and “lightens” if experimental evidence showed that it is necessary to prevent the development of dark pigments or sufficient to lighten pigmentation when mis-expressed (**Supplementary Table S4**). If a gene had been shown to have different effects on pigmentation depending on sex or body segment, we listed its pigmentation role as “context-dependent”. Because the majority of phenotypes we observed in the miRNA overexpression and competitive inhibition screens were seen in the female A6 segment, we also recorded any previously described effects of the gene’s loss-of-function on pigmentation in this segment (**Supplementary Table S4**).

Using the TargetScan 7.2 database (Agarwal et al. 2018), which lists all miRNA 7-mer and 8-mer seed matches in 3’ UTRs of the most-abundant transcript for each gene in the *D. melanogaster* genome, we queried the pigmentation genes on our annotated list, filtering to include only seed match predictions for the 15 miRNAs found most likely to impact pigmentation development in our screens. Because miRNAs typically repress expression of their target genes, predicted miRNA-target pairs where the miRNA overexpression phenotype (darkening or lightening) is the opposite of the pigmentation gene’s previously demonstrated function (darkening or lightening) represent the most promising candidates for biologically-relevant interactions within the pigmentation network.

**Figure 5** shows (a) the pigmentation phenotypes caused by overexpressing each of the 15 miRNAs, (b) the pigmentation promoted by the pigmentation gene, and (c) whether the TargetScan 7.2 database identified a possible interaction between each miRNA-target gene pair. Note that *miR-2b-1* and *miR-2b-2* have the same seed sequence, so are predicted to potentially interact with the same set of pigmentation genes (**Figure 5**). This analysis showed that each of the 15 miRNAs examined has the potential to bind to the 3’ UTR of 3 to 32 pigmentation genes, with an average of 16 potential targets per miRNA. The 24 pigmentation genes that were not predicted to interact with any of the 15 miRNAs were excluded from **Figure 5**.

**Figure 5.**
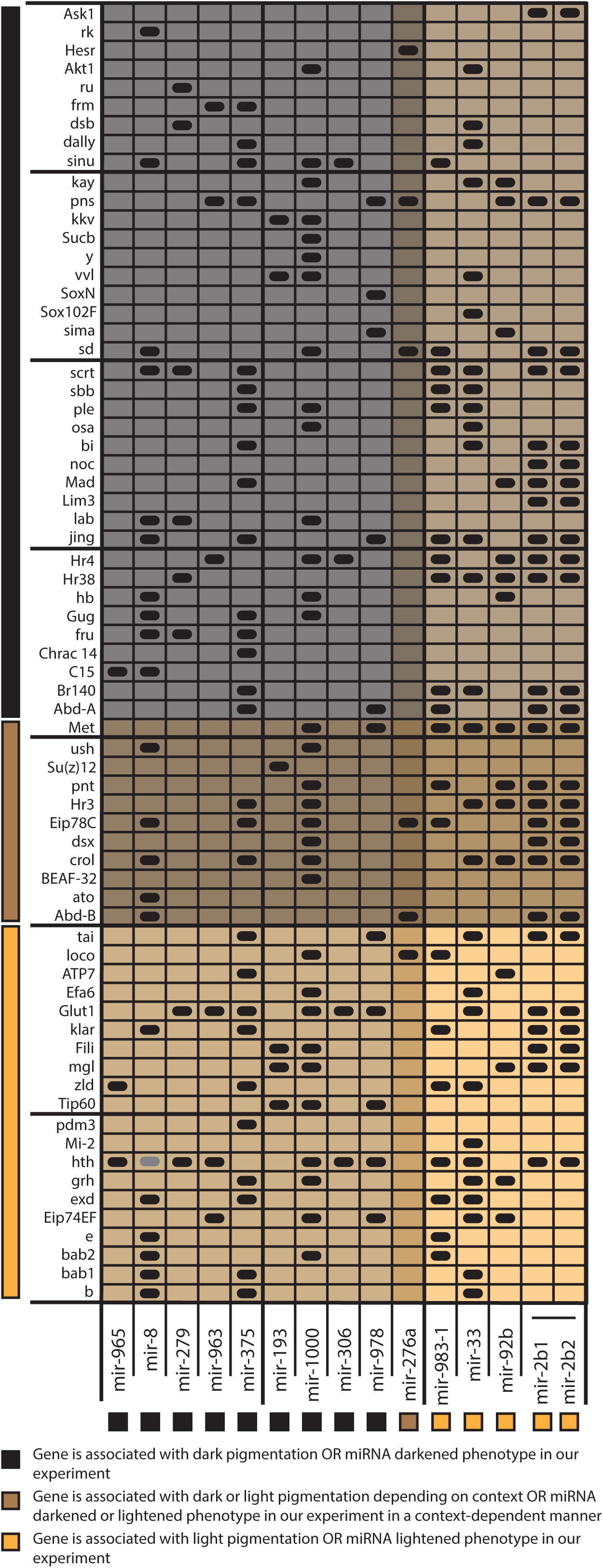
Potential pigmentation gene targets of miRNAs. The table shows miRNA seed sequence targets predicted computationally for the 15 miRNAs that showed reciprocal phenotypes when overexpressed versus competitively inhibited (columns) in the 3’UTR sequences of 69 known pigmentation genes (rows). 24 additional pigmentation genes were also examined but showed no seed sequence matches for any of these 15 miRNAs. *miR-2a1* and *miR-2a2* have identical seed sequences according to the TargetScan 7.2 database, so their predicted targets are identical. The darkest shading indicates the 38 pigmentation genes (rows) and 9 miRNAs (columns) that promote darker pigmentation. The lightest shading indicates the 20 pigmentation genes (rows) and 5 miRNAs (columns) that promote lighter pigmentation. The intermediate shading indicates the 11 pigmentation genes (rows) and 1 miRNA (column) that can lighten or darken pigmentation depending on the context. Cells with black ovals indicate the presence of one or more seed matches to the indicated miRNA within the 3’ UTR of the most abundant transcript isoform of the indicated pigmentation gene. A grey oval was added at the interaction of *hth* and *miR-8* even though this interaction was not predicted by TargetScan 7.2 because manual inspection of the *hth* 3’ UTR sequence identified a *miR-8* binding site (**Supplementary File S1**).

To determine whether potential miRNA-target gene interactions were enriched for cases where the overexpression of the miRNA resulted in the opposite type of pigmentation promoted by the target gene, we used a Fisher’s Exact Test (FET) for each of the 14 miRNAs that only lightened (n = 5) or only darkened (n = 9) pigmentation when overexpressed to compare the proportion of target genes with sequences complementary to the miRNA seed among the 25 potential target genes that promote only lightened pigmentation and the 56 potential target genes that promote only darkened pigmentation. (The 1 miRNA and 12 target genes that have been shown to lighten or darken pigmentation depending on the context were excluded from this analysis.) For all 14 miRNAs, we saw no significant difference in the proportion of possible target genes that lighten or darken pigmentation (p > 0.05 for all 14 miRNAs).

Nonetheless, there are a number of possible miRNA-target gene pairs identified by this analysis that are intriguing based on what is currently known about the development of adult pigmentation in *D. melanogaster*. For example, the *Hormone receptor 4* (*Hr4*) and *Hormone receptor-like in 38* (*Hr38*) genes both promote development of darker abdominal pigmentation (Rogers et al. 2014; Kalay et al. 2016) and were identified as possible targets of 6 of the 7 miRNAs found to lighten abdominal pigmentation when overexpressed (**Table 1**, **Figure 3**, **Figure 5**). This number of predicted targets is a significant enrichment relative to miRNAs whose overexpression darkens pigmentation (Fisher’s Exact Test, p-value for *Hr4* = 0.02, p-value for *Hr38* = 0.002).

For *miR-276a,* which both lightened and darkened pigmentation in different regions of the abdomen, six possible target genes were identified (*loco*, *Abdominal-B* (*Abd-B*), *Ecdysone-induced protein 78C* (*Eip78C*), *scalloped* (*sd*), *pinstripe* (*pns*), and *HES-related* (*Hesr*)) (**Figure 5**). Three of these genes (*loco*, *Abd-B*, and *sd*) have loss-of-function phenotypes that are similar to the *miR-276a* overexpression phenotype: reducing activity of *loco* causes increased pigmentation in female A6 (Dembeck, Huang, Magwire, et al. 2015) (as shown for *miR-276a* overexpression in **Figure 4G’’**), reducing activity of *sd* causes lighter pigmentation in male A2-A6 (Kalay et al. 2016) (as shown for *miR-276a* overexpression in **Figure 4G’**), and reducing activity of *Abd-B* causes lighter pigmentation in A5 and A6 of males (as shown for *miR-276a* overexpression in **Figure 4G’’’**) (Rogers et al. 2014; Kalay et al. 2016). This observation suggests that the complex changes in pigmentation observed when overexpressing *miR-276a* might be due to its repression of multiple pigmentation genes.

Finally, *miR-8*, which darkened pigmentation of the female A6 segment when overexpressed and lightened pigmentation when inhibited (**Figure 3**; Kennell et al. 2012), was found to have a seed sequence complementary to sequences in the 3’ UTRs of four pigmentation genes that promote development of lighter pigmentation in adult flies: *ebony, black, bric-a-brac-1* and *bric-a- brac-2* (**Figure 5**). This observation suggests that *miR-8* might impact pigmentation by coordinately regulating both enzymes (*ebony* and *black*) and transcription factors (*bric-a-brac-*1 and *bric-a-brac-*2) that have similar phenotypic effects.

### miR-8 can repress expression of genes with 3’UTR sequences from four pigmentation genes

To test the hypothesis that *miR-8* affects pigmentation by repressing expression of multiple genes with related functions in pigmentation development, we used cell culture experiments to determine whether overexpression of *miR-8* was able to repress expression of a reporter gene containing the 3’ UTR sequence from pigmentation genes.

As shown in **Figure 6A**, the Black protein promotes the formation of yellow pigments that give the adult cuticle a lighter appearance by catalyzing the conversion of uracil into β-alanine (Wright 1987; Phillips et al. 2005). This β-alanine is then chemically combined with dopamine to produce N-β-alanine in a reaction catalyzed by Ebony (Wright 1987). Expression of the Tan protein, which promotes the formation of dark pigments by catalyzing a reaction converting N-β-alanine back into dopamine (Wright 1987), is directly repressed (transcriptionally) by the Bab1 and Bab2 transcription factor proteins (De Castro et al. 2018). In this way, expression of Bab1 and Bab2 in the posterior abdomen of females, but not males, prevents the extensive dark pigmentation that develops in the A5 and A6 segments of males from forming (Kopp et al. 2000). If *miR-8* normally represses expression of these genes in the posterior abdomen post-transcriptionally, causing darker pigmentation to develop, then inhibiting the activity of *miR-8* should increase expression of Black, Ebony, Bab1, and /or Bab2 proteins in their normal expression domains, leading to development of lighter pigmentation in the posterior abdomen in females, which is the change in pigmentation we observed in our competitive inhibition experiment (**Figure 6B**).

To test whether *miR-8* is able to directly repress expression of *ebony*, *bab1*, or *bab2* by interacting with sequences from their 3’ UTR, we made multiple LacZ reporter genes, each with 3’ UTR sequence from one of these three genes, and tested the ability of *miR-8* to regulate their expression in cell culture. For *ebony*, the reporter genes included the full 3’ UTR sequence, whereas for *bab1* and *bab2*, they contained 683 of 1953 bp and 529 of 1539 bp of their 3’ UTR sequences, respectively. We also made a positive control version of the LacZ reporter gene with a 3’UTR containing two copies of sequence complementary to the *miR-8* seed sequence, and a negative control version in which the reporter gene lacked a 3’ UTR (other than a polyadenylation signal). Finally, reporter genes were made that contained mutations in the predicted *miR-8* binding site of the *ebony*, *bab1*, and *bab2* 3’ UTR sequences tested.

For each of these transgenes, the plasmid containing the reporter gene was transfected into *D. melanogaster* Kc167 cells along with two other plasmids, one driving expression (or not) of *miR-8* and one expressing Luciferase, which served as a control for transfection efficiency (**Figure 6C**). Expression of LacZ, *miR-8*, and Luciferase from these plasmids was controlled by the same constitutive *actin* promoter (**Figure 6C**). Each transfection experiment was performed in duplicate, on at least 3 days, measuring the amount of expression from the reporter gene (LacZ) through Beta-galactosidase activity as well as the amount of Luciferase activity. Beta-galactosidase activity was fitted to a linear model including Luciferase activity as a covariate and both *miR-8* overexpression (presence or absence) and experiment date as main effects. We then compared Beta-galactosidase activity for each LacZ reporter gene between samples with and without overexpression of *miR-8*.

**Figure 6.**
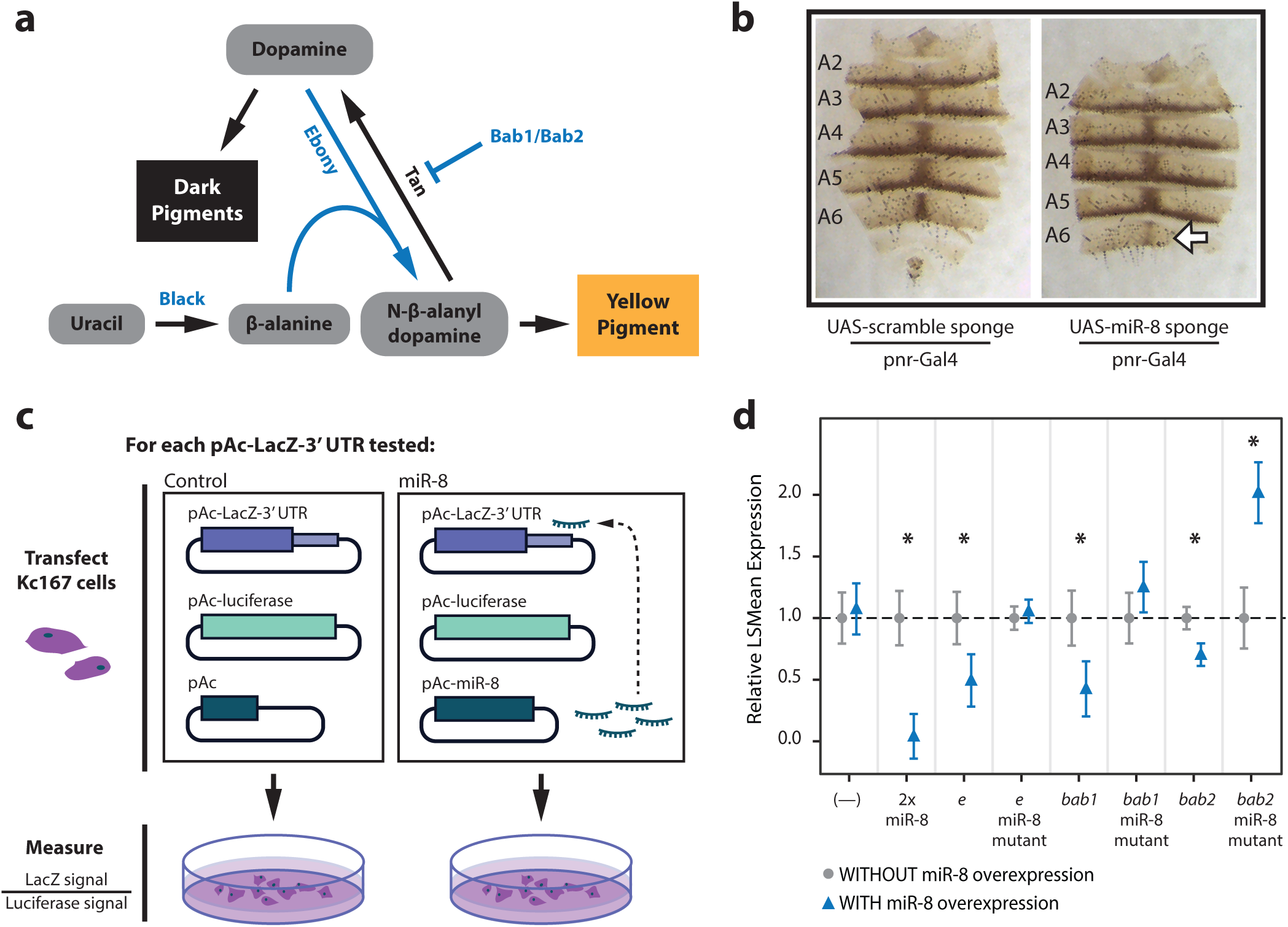
*miR-8* may promote darker pigmentation by repressing expression of *ebony, bric-a-brac1* and *bric-a-brac2*, which promote lighter pigmentation. **(A)** Schematic shows of a portion of the *D. melanogaster* pigment biosynthesis pathway, highlighting the enzymatic functions of the Ebony and Black proteins, which catalyze reactions involving uracil, beta-alanine, and dopamine that are necessary to produce N-beta-alanyldopamine (NBAD). NBAD is a precursor for the yellow pigment found in the cuticles of adult *Drosophila*. The transcription factors Bab1 and Bab2 repress expression of the Tan enzyme (Rebeiz and Williams 2017), which converts NBAD back into dopamine (True et al. 2005), which is then used to produce dark pigments. Genes with *miR-8* seed sites in their 3’ UTRs are shown in blue. Arrowheads represent chemical reactions in the pigment biosynthesis pathway. The flat-headed arrow represents transcriptional repression. (**B)** Dissected dorsal abdominal cuticle from female flies expressing either the control miRNA scrambled sponge (left) or the *miR-8* sponge (right) in the *pnr-Gal4* expression domain. White arrow indicates the lighter pigmentation caused by competitive inhibition of *miR-8*. (**C**) Overview of cell culture experiments used to test for *miR-8* repression of gene expression via 3’UTR sequences from pigmentation genes. The schematic shows that Kc167 cells were transfected with a LacZ gene containing 3’UTR sequence from a pigmentation gene under the control of an actin promoter (pAc-LacZ-3’-UTR), a luciferase gene controlled by the same actin promoter (pAc-luciferase), and either an empty expression plasmid containing only the actin promoter (pAc) or an expression plasmid using the actin promoter to drive expression of *miR-8* (pAc-miR-8). As shown in the right box, the pAc-miR-8 plasmid produces miR-8, which is then available to potentially interact with the 3’UTR sequence attached to the LacZ gene and repress production of LacZ. Cells were then collected from the culture dishes shown, proteins were extracted, and activity of LacZ and Luciferase were measured (not shown). The activity of LacZ was divided by the activity of Luciferase to normalize the LacZ signal for transfection efficiency. (**D**) The least-squares mean (LSmean) of relative LacZ reporter gene expression in Kc167 cells, measured from Beta-galactosidase activity fitted to a linear model including Luciferase as a covariate and *miR-8* overexpression (presence or absence) and experiment day as main effects, is shown for each reporter gene in each condition. In each case, expression was normalized relative to the LSmean for the sample with the same reporter gene without *miR-8* overexpression. Error bars show 95% confidence intervals of the LSmean from the linear model. Blue = samples with *miR-8* overexpression; Gray = samples without *miR-8* overexpression. All transfections were performed in duplicate, on each of at least 3 different days. The X-axis labels describe the 3’ UTR sequences carried by each LacZ reporter gene. “(—)” = a negative control, no added 3’ UTR sequence. “2x miR-8” = a positive control, 3’ UTR composed of two perfect complements to the mature *miR-8* sequence. “e” = full *ebony* 3’ UTR sequence. “e miR-8 mutant” = same as “e”, but with the predicted *miR-8* binding site mutated. “bab1” = a 683bp region of the *bab1* 3’ UTR containing the predicted *miR-8* binding site. “bab1 miR-8 mutant” = same as “*bab1*”, but with the predicted *miR-8* binding site mutated. “bab2” - a 529bp region of the *bab2* 3’ UTR containing the predicted *miR-8* binding site. “bab2 miR-8 mutant” - same as “*bab2*”, but with the predicted *miR-8* binding site mutated. Asterisks show statistical significance (p < 0.05) of comparisons for each reporter gene from t-tests contrasting the LSmeans of samples with and without *miR-8* overexpression. From left to right, p-values for all comparisons were 0.610 (—), <0.001 (2x miR-8), 0.002 (e), 0.471 (e miR-8 mutant), 0.002 (bab1), 0.086 (bab1 miR-8 mutant), <0.001 (bab2), and < 0.001 (bab2 miR-8 mutant).

Compared to the reporter gene lacking a 3’ UTR, we found that expression of the positive control reporter gene with two *miR-8* binding sites in its 3’ UTR was significantly reduced (**Figure 6D**). Expression of the reporter genes with 3’ UTR sequences from *ebony*, *bab1,* and *bab2* were also significantly reduced by overexpressing *miR-8* (**Figure 6D**). Mutating the predicted *miR-8* binding site in the *ebony* 3’ UTR resulted in reporter gene expression equal to the reporter gene lacking a 3’ UTR (**Figure 6D**), suggesting it eliminated all repression attributable to overexpression of *miR-8*. Mutating the predicted *miR-8* binding site in the 3’ UTR sequences from *bab1* and *bab2* also eliminated repression attributable to overexpression of *miR-8*, but unexpectedly caused a *miR-8* dependent increase in reporter gene expression (**Figure 6D**). These unexpected increases in reporter gene expression could be caused by (among other possibilities) the mutation disrupting binding of another miRNA also repressing this reporter gene in Kc167 cells or by the overexpression of *miR-8* changing activity of positive regulators of the mutated version of these reporter genes.

We also tested expression of these reporter genes with 3’UTR sequences from *ebony*, *bab1*, and *bab2* along with additional reporter genes containing 3’ UTR sequences from *black*, *homothorax (hth)*, *pale (encoding Tyrosine Hydroxylase)* and *Hormone receptor-like in 38 (Hr38)* in another type of *Drosophila* cells, S2 cells. We observed the same effects on the *ebony*, *bab1* and *bab2* reporter genes in both S2 and Kc167 cells, and the additional reporter genes tested in S2 cells suggested that *miR-8* might also be able to repress expression through 3’ UTR sequences from *black*, *hth*, and *pale* (**Supplementary Figure S5**). Of these three genes, only *black* was predicted to have a potential binding site for *miR-8* in the TargetScan v7.2 database (**Figure 5**). However, we found that the *hth* 3’ UTR also contains the AGTATTA sequence predicted to bind *miR-8* in the *ebony*, *black*, *bab1*, and *bab2* 3’ UTRs (**Supplementary File S1**). The *pale* 3’UTR did not contain any exact matches to this sequence. Analysis of additional reporter genes, including reporter genes with mutations in potential *miR-8* binding sites in the *black*, *hth*, and *pale* 3’ UTRs, is needed in the future to better understand these potential regulatory relationships. The *miR-8* binding sites identified in the *bab-1*, *bab-2*, *ebony*, *black*, and *hth* 3’ UTRs were all highly conserved among *Drosophila* species (**Supplementary File S1**).

Taken together, these cell culture data show that *miR-8* can repress expression through 3’ UTR sequences of multiple pigmentation genes. It remains to be seen, however, whether *miR-8* interacts with and represses expression of these genes *in vivo*. The Ebony, Bab1, and Bab2 proteins are all known to be expressed in epidermal cells during the pupal stages when adult pigmentation develops (Kopp et al. 2000; Wittkopp et al. 2002), but the spatiotemporal expression of *miR-8* remains unknown at this developmental stage. A prior study testing for physical interactions between miRNAs and target genes in S2 cells did not report evidence of native *miR-8* interacting with *ebony*, *black, bab1,* or *bab2* (Wessels et al. 2019); however, data from Klonaros et al. (2023) indicate that these three genes are very lowly expressed in S2 cells (13, 0, 9, and 32 reads observed from each gene, respectively, out of 17.8 million reads), suggesting that there was little opportunity to identify physical interactions between *miR-8* and these genes in the Wessels et al. (2019) study.

## Conclusions

This study shows that miRNAs help regulate the development of adult body pigmentation in *Drosophila melanogaster*. Through systematic gain-of-function and loss-of-function screens, we identified 15 miRNAs with strong evidence of affecting normal pigmentation development, expanding our understanding of post-transcriptional regulation of this complex trait beyond the three miRNAs (*miR-8*, *miR-193*, and *miR-33*) previously identified. The A6 abdominal segment of females was particularly sensitive to changes in miRNA expression, consistent with prior work showing sensitivity in this segment to genetic variation, temperature, and changes in the expression of other genes. We found that many pigmentation genes contain predicted binding sites for these 15 miRNAs in their 3’ UTRs sequences, suggesting that these miRNAs might impact pigmentation by directly regulating expression of these genes. Indeed, our cell culture experiments confirmed that *miR-8* can repress gene expression through 3’ UTR sequences from at least three pigmentation genes (*ebony*, *bab1*, and *bab2*), all of which promote lighter pigmentation.

Our findings support the emerging view that miRNAs influence trait development by coordinately regulating expression of multiple genes with related functions. For example, *miR-8* has the potential to repress expression of both enzymes (e.g., Ebony and Black) and transcription factors (e.g., Bab1 and Bab2) required for proper pigmentation development. Similar effects have been observed for *miR-iab-4* and *miR-iab-8*, which are encoded in a single transcript within the *D. melanogaster* Hox locus. These miRNAs have been shown to coordinately regulate the Hox genes *Abd-A* and *Ubx* as well as the Hox cofactors *exd* and *hth* in the larval ventral nerve cord in a manner that is essential for proper segment patterning, fertility, and mating behavior (Garaulet et al. 2014; Garaulet and Lai 2015). Other examples include *miR-196* regulating multiple Hox genes involved in limb patterning (Hornstein et al. 2005), individual miRNAs regulating batteries of genes involved in synapse formation and neuronal identity (Barca-Mayo and De Pietri Tonelli 2014), and miRNAs coordinating expression of cell cycle regulators to control organ size (Brennecke et al. 2003; Hyun et al. 2009; Verma and Cohen 2015). The miRNA-mediated fine-tuning of gene expression networks appears to be a conserved mechanism for achieving robust and precise developmental outcomes (Tsang et al. 2007).

For the last 20 years, *Drosophila* pigmentation has been a powerful model system for understanding both developmental and evolutionary mechanisms at the molecular level. This study shows that it is also well-suited for studying miRNA-mediated regulation of gene expression and phenotypic development. Future work determining which predicted miRNA-target interactions occur in vivo during pigmentation development, characterizing the spatiotemporal expression patterns of these miRNAs, and testing whether natural variation in these miRNAs contributes to pigmentation differences within or between species will further illuminate the roles of post-transcriptional regulation in trait development and evolution. Lessons learned from studying this trait may also apply to other complex traits in *Drosophila* and other species.

## Data availability

As noted above, raw data for cell culture experiments and the R code used to analyze them are available at 10.5281/zenodo.20451261. This Zenodo record also includes >1200 images of flies collected as part of the overexpression and competitive inhibition screens. Sequences of 3’ UTRs used in the cell culture experiments are included in **Supplementary File S1**.

## Acknowledgements

We thank members of the Wittkopp lab, including Ayse Tenger-Trolander, Elizabeth Walker, Alisha John, Lisa Kim, Alicia Wang, Nastya Vlasava, Anati Azhar, Tasmine Clement, Patricia Simmer, and Shibo Hou for advice and assistance with this work. We are also grateful to Laura Buttitta for providing access to equipment and expertise required for the cell culture experiments and to Scott Pletcher for sharing a double balancer fly strain.

