## Supplementary Figure and Table Legends for "microRNAs affecting development of body pigmentation in adult Drosophila melanogaster"

**Supplementary Table S1. Summary of data from overexpression and competitive inhibition screens.** This table includes identification numbers for fly strains, miRNA annotation confidence, counts of flies collected from each genotype, statistical tests of viability, brief descriptions of pigmentation phenotypes, and brief descriptions of non-pigmentation phenotypes/defects observed. The first sheet (Table_S1_summary_of_screens) contains all the data. The second sheet (Column descriptions) contains definitions of the column headings in the first sheet.

**Supplementary Table S2: Phenotypes and notes on the 7066 flies scored in miRNA overexpression screen (including controls). Sheet 1 (All_OE_data):** Each row contains the gene under UAS control (“UAS_line”) for each cross, the miRNA associated with the line, date of collection, date of phenotyping, sex, balancers present, notes on phenotypes observed, classification of phenotype in posterior abdomen, anterior abdomen, and thorax (“l” for lightened, “d” for darkened, blank if unaffected), presence of visible developmental defects in abdomen, thorax, bristles, or wings (1 = defects observed), whether the defects obscure the pigmentation phenotype (1= slightly obscured, 2= completely obscured, 0=not obscured), and whether observed defects occurred in the same body segment as the observed pigmentation effects or different body segment (“s” = same, “d” = different). **Sheet 2 (Summary_data):** Summary phenotype results of miRNA overexpression crosses. The headers “T”, “AA” and “PA” refer to the thorax, anterior abdomen, and posterior abdomen, respectively. Phenotype scoring: “L” = lightened in the *pnr-Gal4* expression domain, “D” = darkened in the *pnr-Gal4* expression domain. Only phenotypes observed at a penetrance of at least 50% are marked. “n” refers to the number of pnr-Gal4/UAS-miRNA individuals phenotyped for each cross. Viability was recorded for each cross, where a designation of “reduced” viability indicates a significant result from a one-sided binomial test comparing the proportion of pnr-Gal4/UAS-miRNA individuals collected relative to those inheriting balancers in place of pnr-Gal4 or UAS-miRNA. Crosses that produced 2 or fewer individuals are marked "Lethal." This summary table is an expansion of **Table 1** to include all 166 miRNAs tested in the overexpression screen.

**Supplementary Table S3: Phenotypes and notes on all 3745 flies collected in miRNA competitive inhibition screen.** Each row contains the miRNA sponge or control (scrambled) sponge under UAS control ("UAS_line") for each cross, the miRNA(s) associated with the line, the rearing temperature, date of collection, date of phenotyping, sex, balancers present, copy number of sponge transgenes inherited (default 2 copies, one on each of 2nd and 3rd chromosome, "1" indicates single copy of sponge transgene), notes on pigmentation, phenotypes observed, classification of phenotype in posterior abdomen, anterior abdomen, and thorax ("l" for lightened, "d" for darkened, blank if unaffected), notes on non-pigement defects or other phenotypes, presence of visible developmental defects in abdomen, thorax, bristles, or wings (1 = defects observed), whether the defects obscure the pigmentation phenotype (1= slightly obscured, 2= completely obscured, 0=not obscured), and whether observed defects occurred in the same body segment as the observed pigmentation effects or different body segment ("s" = same, "d" = different).

**Supplementary Table S4: List of experimentally verified pigmentation genes**. For each gene, the table includes name, annotation symbol ("CG" number), categorization as transcription factor (Y = annotated as transcription factor, N = not annotated as transcription factor), pigmentation role ("darkens" if gene has been experimentally demonstrated as either necessary or sufficient for the development of dark pigmentation; "lightens" if experimental evidence shows that gene is necessary to prevent the development of dark pigments and/or sufficient to lighten pigmentation where misexpressed), loss-of-function effect on female A6 melanization, and supporting citations.

Citations: 1 = Kalay et al. (2016), 2 = Rogers et al. (2014), 3 = Dembeck, Huang, Magwire, et al. (2015), 4 = Wright (1987), 5 = Wittkopp et al. (2003), 6 = Kopp and Duncan (1997), 7 = Riedel et al.( 2011), 8 = Dewey et al. (2004), 9 = Wakabayashi-Ito et al. (2011), 10 = Norgate et al. (2006), 11 = Shakhmantsir et al. (2014), 12 = Baker and Truman (2002), 13 = Sekine et al. (2011)

**Supplementary File S1. 3' UTR sequences attached to the LacZ reporter gene for testing inhibition by *miR-8* in cell culture.** This file contains sequences of 3' UTRs tested in cell culture, annotated to show sites expected to be bound by *miR-8* and with information about the conservation of these putative *miR-8* binding sites in other *Drosophila* species. Sequences of mutated binding sites are also included.
