## Supplementary File S1 for "microRNAs affecting development of body pigmentation in adult Drosophila melanogaster"

**Supplementary File S1. 3’ UTR sequences attached to the LacZ reporter gene for testing inhibition by miR-8 in cell culture.**

>***bab1* 3’ UTR** – 1953 bp; *miR-8* binding site (bold) is perfectly conserved in 21 of the 22 *Drosophila* species examined; the sole exception is *D. mojavensis.* In the mutated version of this UTR, the bolded sequence AGTATTA was changed to CTGCGGC.

AGGGAGTAAAGGGAGGGTGAAACGAAGGAAATGATAAAGTTGAGAAATGATAATGGGTGAATGAACGCAAATCAGAAGCTTCGGCAGCTTTACTTGGCCTGTGTAAGCCTACGCCTTGGTTAAATTAAATTAATTTAAATTATATATTATTTATTATACGTGGAATCTGTAAACTGCATTCCCCGATTTATGCCCATAGTGCAAATTGATTGAATTTGGCCCGAGAAAACCGCATTAGGCATTGCATCCACATATTGTAAACGAGTTACTTTCCAACTACATGATATATCACATTTCAAATTATAATTATGTGTAAATAAACTAGATATATACGTTTTGAAGGTTTCCTCCAAATCTTTGCGTTTTCACCTCGAACAATTCCTACACTTGATATCCAATCGAATTTACAAATAAGCTTTAACCACGTTCTGTTTATCATATCAAAAGTGTACATTGCAATCTATTGTAGTTAACCAAGGTGTAGGCTGTCGCATTAGCCAGTTTTACTCTAGTTTCCTCCACGCGATCTTCGCCACCTAGTTTCTATTCCTAGCGTACTTACTGAATCAAAATTAA**AGTATTA**TTATGTTATAGTCCATCAGTCTAACTAACTAACTAAATAGCGACCGATCTCTTTAGTTGTAAGCTACAGCGAAGGTGGAAGCAGCAAACCCGGAACATTCCACAAACCATTCGAAATCTGTTTCGAATGCAACTTCAAACTTCAATTGAAACCGAATCAGAAGCAGGATGATCCACAAATCAAAGCAGTTTAATATTAATGAAATGGAATAAAATCAAAAGAGTTTAAGCGGATTTAACTTGGTAATTGTGTGCCCGAATTTGGTTTAGTCTTTAGTTCACGAGAAGCGAGAAAGAAAATGTACAAAATTATCTAAAAATGAAAAAAAAAAAAATAAAAATTGAAATCGAAGCTTGTGGCTTCGAACGCAAATGGAATCAACTAAATTTAATGTTTAGGCTTAACAAAAAACAAAGAAATTCACAAAAAATGCCTTTCGGAATTAAATTCAGCGATGAATACACCGCAAACAAAATTTGTAAAAAGAAAACAAACAAAAACACGTGAACTTTTTTAACGTATTTATACAATCGAAATTTTTAGACAAAATTTAGAAGCAACAATTTTTTACACCGAGCCCTGCTGCTGCTCCCGCCGAACTCGAGGCCACTGATTTCGACATTATTGGAAACCCAACGTAGTCCCCGAATTGATCCCGCATCAACCCAAATCCCCACTAGATCCCCTTCTAAAGTGCCTTTTCTAGCCTTAATTTGGACTCTGCTCCACTTCCGATCCGGTCGAGAGCGAGGCCAAGGATTCGCGGCAGGACTCGAGGATTTATTTGGCAAGTCTGGCGACATTACGATACGATTGAGTATCGGAGAACGTACATATATGAGTATCTCTTTATTTGGTTTAAACCCCTTTTAGGCATTTAGGCATACTTTCGGCAACGTTTGGTTTGGCAAGCACACAATTGATTAATCGGACAAGAGTTAATTACGATGATAATGATAAACATGACTATTTTTGCATACGCAAGAACATCTACCATAGCCATAGCCCGTATATAAGCACACACGATATATAAGTATCTATGTAATCCCAGTTCCCCCAATAGACTTTGATGTTCCCACACGTATCGAAACTTAACGAATTACTTTCGGAATAGGATGTAATATAAGCAGTGGCAAGAGTTAGCGGCTAAATCTCCTGGATATTGTGTATCTTTGTACGATCTGGATCAGCGTCCAAGTCCACGTCCAAAACTCAACATTTTCTATAAACTTTAGTGCAAACTTTATGTTCAAATTAACTTTAAATGTCGAAATGAATGTTCTAATTAATCGAAGAAAACAAAAAACATAAAAACTTAAAAACGGAAAACCGAAACGAACAAAAAGTATTTTATAAAAACATTGAAAAGT

>***bab2* 3’ UTR** 1539 bp; *miR-8* binding site (bold) is perfectly conserved in the 22 *Drosophila* species examined. In the mutated version of this UTR, the bolded sequence AGTATTA was changed to CTGCGGC.

ATCCTGGGCAGGGGCGTGGCATTGGGACACACGCTGGGGTGGGGCAATCATTAGGTAACCAAATAAATGCGAGGGGTGCTAAGTAATCTTGGATAAGAGTAAGTAGTTTTGATTTTAGGACTGGAAGGTCTAGCAATTTCAATTTTATAATGTGCAAGACATATATTTCTTAGAGCAGCAAGATACATAGCATTACACATACGGTTTTTTTTTTTATGATAAATCTTTTAAAATAGGAAAAGTGTGTAGGAAGCCGCACAGCTCATATAAAATGTGACTGATAACGTAAAACACACGTTATATCTTATCAAGAAATAGCAAAAACGTGGAATTAAATGAAACACAAATTCACACCAATGACATCAATATACCACGTCAATACCAAGCATATGCGATTTATTTTCCCCAAGAAATAGCCCGAAAAGTCCAATGATTACTTGCCCCAAAGATTCTTGGCCCCAATTGGATAACCACCCTGAGCCCAAAGGAAGAAAACAATAGAAGACCTACCAACAAACAAACATATTTGGACTGCTACTTGAACCCCAACGTTTTTAGTGTTTAGTGCTTAGTTGAAGTTGCTAACTGGTTTTTATGCCTAGGCTAGTTTTAGAGTGTATCATCGAGGAAGGTATAA**AGTATTA**ACCTATTATTAAAGCACACCCCAAAACACAACAAACACACACCCACACACATACTTCGGAACTTTCCTGTAAGGATAGGATCTCGTACTTTGTCTTATTTATTTCGAGAAAATTCGTTGCGTTGATTGCCTTTTTTGCCACGCGAAAATAATAATAAATAGGCTTACGATTAACAATAAACAACGTAACCGTAAACATTAAAGTTACGATTAGAGACTTGTGCATTTCTTAAATGTAGCTTAACCCTAGAATAGCCCTCAAATACAAAACACTGAATGTTGTCAAATCCGGTAATCACCACTTTATACATACATATATATATATACGAGTACATACATATGTATGTACATATATACGCAATACTTTAGATACATAAACCCATACAGATAGACGTGCATACCGTACCACGCACGTACATAAACTTATATTAATGGTATTTCCAAATGTAACTGTAAATAATGTACAAATTTCCATAGCTTTAGGCTTCGACTTAATCTTAGACTTACCCCCAAAAAGCCCTTCGCCGCACTTCTTCTATGGTAATCGTATTGTAAATGAACGAATGGGCAAACAACGCAGGCAGCATCACATTAAATTTGTGCTAATTTTATGTAAACAACCACACAGACACAAAGTCACACAGATTACACAATACCATACCACCACATGGCACACACACTCGACTGTCAGATACGCATATGTGTGTATATGTGCATATGTATTGTCCGAACTGCGTGGGTGAAAACTATTATTATTATAACTATTATTACGATTATTATTATTACCTAACCCTAGAGATGCGCGAGGCATTCTTGAGTTTAGTTAAACCGTAAATTATACAACAATAAACAAAGAAAGAGAAAATAAAAACGAATCGAAAAGCTAAATGTCAACC

>**ebony 3’UTR** – 203 bp; *miR-8* binding site (bold) is perfectly conserved in the 19 *Drosophila* examined. In the mutated version of this UTR, the bolded sequence AGTATTA was changed to CTGCGGC.

GACGACCACCCGGTGGACGTGTCCCCAAACCAGATCTGCAGCGAATTTATCCCGCACCACGTATTATATAGTTAACTAGCATAACCTACCACCT**AGTATTA**GCCCTAGGCATAGTGTAACTTAGCGAGAAGTTTCGTAAGTTTGTGTTGACTAGTTGTAAGCCTTTGATTTCCAGACAATTAAATACACAACAATCAATAATT

>***black* 3’ UTR** – 268 bp; *miR-8* binding site (bold) perfectly conserved in 18 of 20 other *Drosophila* species examined; exceptions are *D. virilis* and *D. grimshawi*, which are the two most divergent species from *D. melanogaster* among the 20 species examined

ACGGTGACGCGGCGATGGCATGTTATGAGGTTAGGTGTTGACAACATTCTCTTTAGGCTTACGCCGATTGTATCTTCTAGTGTATGGTTAACTAATACTACTCACGTTTGTATATTTTTCTGGATGTTA**AGTATTA**CCAAAAACAAAGAACAAAAATACATATTTATAGGTCATTTCATTTAGATTAAAGTGTGTCCTTGCCTTGGGAACTGGGAAACGTATCGGCAACAGGATGCTTTAAATGCAAGGTTATTTAAAAACATAGTGA

>***hth* 3’UTR** – 3816 bp; *miR-8* binding site (bold) was not identified in TargetScan, but it is a perfect match to the sequence predicted to bind *miR-8* in *ebony*, *black*, *bab1*, and *bab2*. It is conserved in all10 *Drosophila* species examined.

TCCTCGTAATCCGGCCCCGACCCGGAACCGGAAATGAAACTGGAACTGGAACCAGAGAGCTCTCCGCCGGAAGCGATCAAGCCAAAATCATAGCTTAAATTTTACCACCACCCAACACTGGAGACGAACGACGAAGGAAGAAGTACGAAGGAGCTGAATACACATGGATCCGATCCCAAAACTAAAGCATACCGCGTGGCAATATCAGTCAAGTAATAGCAAACAAAAAACAACAAGCAAAGAACAACAATCGTAGTAAAATCTATACAAAATGATATTAGCTAAGCAACAGCGACAAAAACAACAATAACAACAAAAGAAACAGCAGAATCCAGAACAACAAAAAGCACCAGCAACAACAAACCACATCAAAAAGAAACCCAGAAATTTTAAAAGAAACTATTTGTGTATACCCTAAACGTAAATAACAAAAGCAGAAAAAAAAGGGAAAATAAAAGCAAGAAAAAAGAAATATTTATGCATAAACAAAAATGAAACGCTTAGCTATAAGTTAAATAAACAAAACAAAACCAGAACTCACACACACACAGACACACCGAGACGCAACACAACAATTTGAAATGCAAATTTTTATGTTCAAAGCTGCAGAGTGCAAAACACAACATTTTAAAGCATGAATTTAGGCGTAGTCCAGAAAGGATACAAAATTGAAAAGGATATACAAAATTTAAGCGACATTTCGTTCGGATTTGTTATAGATTTTAGTTTGAATTTAGGAAATGCCCCGCCCCACCAGAAAATCAAGGGAAAACCAAACCAATAGCAGCAGCGAAATTCGATCAAGTTTGAATATAATTTGCGCAATGGAAGGACGAGAGGAGAATTCTTTGCTAGTTCTTAGTTTCATTTTAGTTATGTAACAAGTCAGTTGTAGTAACGCTTTGTTTTCTTTACCTTTACGCATATGTATATGTACATACAAAACAAATCGGTAACAAACTATAGATACATACGTACATCTATGCTTAGGCGATAGCACCCTACTATATCCTACGTTTAGACGATTAATTTAAATAAATTGAATTGTTTTCTAATTTTATGCCTTGCAACACTTAAGTAAAAACGAAATGGAATTTAATTTTTACAAAAGGGAAAGATGATAACTAGAGGAGGAACATCATATGATTTGAATTTGTTTTAGAGTTTTCATTGTGTGCGGATCTTGT**AGTATTA**TTTTCCAACTGAAGCACATGGACCGATTGAATATATATATTAATATGTATATGTATGGATTTTATGCGAGATACAAACAATACGATACAAATAATGGCATAGGCATATACATAACTCTGAATATGTATACGTAATATTAATAATAAATTATGCTTACGTTTAAACTGTACTTTTATTTATGTATTCTAATTTGTATGCGTTAATCCTTAGTTAACTATTATTATTATTATTACATTTACTTAATTAATGATTTAAATTTAACTAATTCAAGAGGATAGAATTGTACGAGAGATGATTTATTGATGAAAACAAAATGCATATATGAGAAAATATTTGAAAAAATAATCTTTCTTAGTGTTTTTATTTCAATTTTGGTTCTGATTGGTTATGATTTCTTTTCAGCGCGGGCAAACCCTATTATTTTTGGGGGATATAGACTCCAAGAGCCTTATATCCTCTCCACCCAGGATATCTTTATTTTAATTCGTGTTGGATCTTTTGGCAGACACGTCACATGCTAAGGAACTGAATTGAATTTACATTGCTTGTGAATCTGATTAATCTTAAATGGAAACGATATACGTAAATGCAGAGGGCAGGAAATCGAAATTTAAGTATTTAGTTTGAAACTTATTAACTATGAATAGATTGAAGGTGTGGAAAGCGAGTACCTATTTTGAAAATACAAACGCGAAATAAGATGGATTACTTAGCCTGCTCTTTGAAACTTGAAACTTGAGCTTCTTTTGCATGGGAACCCAACATTATATTGGCAGACACACTTAGGCAAACTCATCTTACTTCTTACCGACTACACTCGAAACAATCACTCAAATGTATACGAACTTCTCTAAGAATCCCCAAGGATGAAAACCTTATGCGAATTTTTTGCACCTAACAAGCGAAGCGATAAATTTATTTACAAAATTTTGCCTAATTGCAAATGCTGTATTTGCGAATAGAAATTGTAATTGAATTGAAGGTAAATCGAACATTTTCACAATTATACATCAAATCTATACGATATACGAGTACGAGTACGCATCGCTAGTAATTTATTAGACTATATATATATAAAGTAGTTAAATAATCGACCCAACAATTGACTTGAGAAACGTTTAAACGCAAATTGTACAGAACGCCTTAAATGAGAAACCAAAGTGGAAGCATTGAATGAATGAATGATGTATACGAGTCTATATACAGAACATAAACTATACCTAGTTATACCTAACTAAATTTTAATGAATCTCGAATAGTATATATATTACTATATATCATACACAACTTATGGCTTAAGAAACGGGAACCAAATGCAGTCGAGTCGGAAGGCAGTGAGTGATATCCCACATTTGTTGGATCCGAGAATTCCCGAGAGTTTCTTAGTATTTTGTATGAGCTTCTTTTATATGCCCCCTCAACTTTCGATCGTGGGGATGTTCAGTATCTGATCCAGACTATTGTATCTGGGGGATATGCTCGCTGCGCCGAACCGACAAAAGCTTGTAAATTAAATCAGTGCGATTGATTATGAGGGTATATGGCAATATTTGTGCGTATCGGATCGATGTGGATAAGCACCCACAATAATCCAGATAATGAGCTATACGATAAATAATACGCATTGTATATGGAAAAGACGAACTCGAAGGCTTTTTTGTTTGAAATCATTTTCGTGTCAGAATTTAAATTAAAATGTGATTTTTCTCGAAACTTTAATGTATACGCCGTAAATATTATGAATATGATTATAAAGTGCTGCAGCAACATTATTATTACCATTATGATTATTACGATTGATAAATCGATTATAAATTTACATATTCAAGTGTAATGTATGTAATAAATATTGTACATTTTTGATAACTTCCCCGTAGACTACACTGTAAATATATCGCAACATTTGCATACATCACCCACACTTAAGTACATTACACTAATTGTAGAGTTAAGTTTAAATCTAATCTAAAGCTAGATCCTAGCGATAAATGGTAAGAGACGGATAGGGAGCAGAAATGCAACCCTGAGCATAGACTAAAATGCACCAGAAAAAAATTAACAAAGGACTTGTTAAAATGGAAGAACAATTTATACAACTTACCCACGCGACAACATCACGTAAATCCAAAAGAATGAAAGAGCTAGAAATAGATAAAATCAACACACTTGCGAGAAAGAGAGAGTTGGCGAGCGAGCGAGCGAGCGACACAGAGCACTAGAGTTAGAAAGGGAGAGCTAACTTAGTAGTGCGAAATCCAGAAATTAAATCAATAAAACAAAAGAAAACTCTAAGAAGTAAAAACTATAAATATATTTATATGGAGAATTGAATTAAATAAATGTTTGTGCGTGTAAAAATTAACTTTAACTAAATTGCTAGACTTAAAAGCCAGAATGAATTAAATATAAATATATTGAAAGGAAACAGTATCATTAAATTAATTAACTAATTGAAGCTTCTATGTAAAGAACTAAAAACCAATTAACGAATTCCTGAACCTGTTGGCGCCACAAATCGCCAGGTAAATGGCGTAGGAACAACAAGGCCGAAACAGAATATGGGGAAATATTGAACATAATTTTCAACTGTAATAAATTATGAAATAAAAGATTATGAAAACC

>***pale* 3’ UTR** – 1614 bp – no *miR-8* binding site predicted by TargetScan, yet repression was seen in cell culture with miR-8 overexpression. No match to the AGTATTA sequence.

GTGGATGGGGGGGAGATATATGTATGATATATAGAAGCACCCAAAAACATCGCCCAAAAAAACACCAGCCACCCCCACCACCCAAACATACACATACAGGTGCAACGGTCTCATTTATACAAATTAATTTTAAATTGACATACCGCACAATCCCTCGTACACGTAATCCTTAAACCAAAATTAAGGCATCGCGCATTATTTTTTATTTTTTGTTTATTGTTTTTTGTTTAATTTTTATTTAATTGTCATCGAAAAGTTGTAATTGTCGAATCGCGCACATGGTAAAAAACACAAAAATCAAGCATATTTATTTGTTGATCACCCGATACACCACCCAACCTGCATATTTATCACCGCAACTATTATTGGACCAAAAAATCTTTAGACGTTTAAGTGATGTGAAAACAAAGCCAAGCCAAGCCAAGCAAAGCAAAGCGAATTAAGTGAAGTGAAAAGCTGGGAAAGCAAAGTTCAGATACATCCGCACTTAGTAGGCAAAGTACCACACAAAAACCGATTGTATACTCAGACAAAAATTGTTTTTGTATTATTATTATCTTCATTTTGCCTCTCACAATATTTACCTAGACTATATAGACACGCGATATAGTATTCCCGGTATAGGAAGCTTCATTTAGAGAGAGTATTTCTAAACAAAAATGATATAGATACAACAATCGAGCAAGGAACAAATGGAAAACAAACAAAAAAAAAAACGGAAAAGATAAAAAATGAAATGGAAAACAAAACAGAATTTTCTTACATTGTACAATAACTGTAAACGACAATATTACTATTGTGTACTATTATTATTATTATCTATTATTATACTTATGACTATTACGTATTGTATTGTAGGTATGTAGAGTTTAAACATTATTATTTTATTTAAGTTTTGTGTATTAATAAAATTAAAAATGCAATTGAAAAATAATTACTCTTTGGTTTTTTCATTTGATTATCTTCTTTGGCAATTCGAAACTCTAAAGTCTGAATCTGTTTAGGGAACATTTTATGACATCAAAATGGAATATTTATTTAAAATTAATGAGTAGAATCGAGGAATATGATAAATATATTTATGGGTGAGAAGAAGTGAATCTCATGTGAACTAAAATCGAGCTATGTAGAAAATATTTCACAAATTTGATGAAATTATATAAAATAACGAAAAACCTTGGACCAATACATATGTATTATTATAAAGTAATTAAAATCAAAATATTTCAAAGTAAACAAAGTAAAACATTGGGATCAAGAATATTACATATTTATAGTTAACGTAGCATGGAATTTTATTGTTATTGTTCGATGTATTGTAGTTGCAATTGGATCGAAAGCCAACCAAGTGTTAACATTTCTGCAAGCAATTAGCCAAACTAATTACTAATTATATTCGCACATAAATCTATATATATAATCAACTTAAAGCGCAGTTGGTCCCCAAGCCAGGCTTCGTTGGTTCTGAAAAATTCAATAAAATTATATGTTTGAATGGAGCACAAAACGTGCAGATATCGTAAATATATATATAAATATATATATATACACATTCTTCTCTATATGTACATTTAATCGCTATTATATGAATAAAGTTGAAATCGTTTAAACTTG

>***Hr38* 3’ UTR** – 3348 bp- no *miR-8* binding site predicted by TargetScan; no match to AGTATTA; no repression seen in cell culture.

AGGCGATCATCAAGCGTATCATCACAACTTGCTTCCTTAAACTAGCCCCTAAGTTATGCCTCCTAGGATATACAGAGAAAGGACCCCATAGGACGGACGCAACTAGCTTTAGTAGAACCCTGAAATAAATAAATCTCACAACAGCAAAAACAAAACCGAACCGAACAGAAATGAAGCGAATAGCAGACCCAGGCCATATCTTTAGTGTAGAGCTAGGTAGTTAGCCGGACAGCCCCGGCTCCTTCGATAATTACGGACATGCATATTTGAGAGGGGGTTTCCAGTGCACAGCCTATGGCTCCTGCGTGACTCGTCAGCACCGCGAGCTCCAACTTGTTGACGTTAATTGTTAAATTGTTTAATTTCAACTGTCAAAACCGGAATCAACGGCCGGGCACGCAATGGCAACACTTTCTATCCCCGGACTTCGAAGCCTGCTCAACATTCGGCACTACGGACGGACAAACAACGGACAGAAACAGAACTCACTCTTGCTCTCTTGCCTTTTGCTAACTTCTAGTCAATTGATTTAGGCGAATCAAATAAATAAATAAATAAAATAAGGGCGTGCAGCAGTAGTGTTATATAATTTCTATGCCAGACCCCAGCGGTTCTCTTCAAGGAAATCCCCCAATGAGTTGCACAAATTGGGATAAAGTACGATAGCCTATTATTCTTATATTTCTTTTAAAAGCTCGAAGATAGATGAGAACTGTGTGGAAATCCACTATCATATCATATAGTTGCTATAAGCCGTGCTTGCCCTAAGCTAAGTTAGACCCGCATAAAGTTGATAGCCCAACCAAGTATTTCGGTTATTTCCTAGACTAAGGTCCTAATAGTTATAGGCTAAGACTATTCTGTTCGATTTATCAATGCACCAAACAGTGCACAATGAGAGTATAAGTACCTTCTTGTGATGATTGTGTCTGACACAGAGAGAGTTGCACACAAGCACACAAACTAGCCGATAAGTTACTAAATACGATCTAATATCTAATATATATAATATAATATAATATATATAAGTCCAAGTATTCGGAAATCCAAGAACCCTTGCATAACCGCAGTTCGTACGTTCCAAACGAGAAAAGAACTTTATTTAATCCTAGACCACTCCATCTAAGTTCTCAAAGAATCGTATGTGGATCGTTGGATCTGTCTCTCTATATATGTGTGTGTGTTATCTCGATAGAAAACCCCTCTATGTGATTTTGTGATAGATTGGCATTGAACTCTATATATTTATATATATATGTCTATAATATATATACACGCATAAATATATATTTTTATGTCTAACTTTTGTATGGTTTATTTTATACGTACCACTTTTCTTTGATAACAAAAAGTAAAAAACTCGTTAGATAGCAAATATTTCAAAGGTATGTTACGAGGACTTTTCAAAGTACCAGTCTTTAGCGACTTTCCAATTAACGTTCGTATTAACGAAAGACAGATTTTCTATGTGTTAAATTGAAGACTTCTATAACTATAACTAAATGCAAGCTAAGAGCAAAAACACAAATCCACAAATCCCCAAAGTGAATAACATATCTCTTCAAGCTTTCGAGTGCACGGAACACGTAGAACCGAAACCCAAGTGTTACTAAATCCATTTAATAATCGGCAAGCCGGGGGCGTCGGCGTGGTTAATACGTTCTCATTACCTATACAATTTAGATAGATCATTATTAAATTATTGTACATGTAGCACATGAAATGTTCGACAACTAGATTTTGTACCATCTTAAAGAAGAACCTAGGCCAAGCTAAACTAAGTATAAACTATGATCTGCATGCGGCTGAGCTGTAGCTATGAGAAATATACCTGCGTGGATCTAAGTGAAATGGGACACTTTGAATTTAGATATGAAACGTTCTAAACGCGACGTACTAACTCTCCCAACTGCGAACTCTACCAATTAAGAGAAATTCCCAGAAAATGTGTCAGGATTTCAAAGCGTCCCATCTCACTTGAACCCACCCAATCAACAAATACAAATCCTAGGGAAGTTGAGAGGTTCAGCAACCATAGAGCAATATTTCATAAGAAAACGCACCTTAAATTACCGAAAAACATAGATTAACCTGATCTTGTAACGTTTGGGAGCGATAATAAGCCAGGATTAAACAGGAACAGTTAGGTGACCAAATCAGTTCGAAACGAGATGATAGATAGGTTCGGGTTCGAAACCCTAAACGCGATGCCATTTTAGCCGTTACAACATTGGATATCAACCATGCACATGAATATGAATATGAATATGAATATTATAGAGATATATCTAGCTATAGGAACCTACTTTGTACCTACACGACATGGAAACATCAAACCTACATGCATATTTACACACATATATTTTGAATAGAGCGACGACTTTTACAAGTTGCGTACAAAGCTATAGCTATAGCTTGATATGGCCATCCCAGAGCGAGCATATACATATATTTTGGGTTATTGTTCTTTTGTAATTTTATAAATGCATACATATTTATTGTACTACGTGAATGTCAAGTGTGGATTCATATTTTTGAGATACAGCTACAAAACGAAACAAAAGAAAATAAAACAAAACAGAAGAGTAAACGTGAAATTTTTCGATGAAACAATTTTAAATGAGAACTTTTTAATATTGCTATTAAAGGATATACATATACACACTAACATACATATATATTTTACTATGTAACGGATAGAATTAAGCTAGATGCAGCGCATAAAGCTTTATACAACAAATTGAAAAGCAACAGAAGAAATTGGCACAAATTAAATTTATATAGCATAATTAGACGTCCTTCGCAAGATAATGTTATTCGTAATAAGAGCGTCAATCGGTACATCGGGCGCTATTTCCCACTACACCCCCAACCACACAATAGATAACCTAAGCTATGTATGTACATTAGCTATGTATATCCAGCCCACTTATGCGCCTACTACTAGAAATGCAGAAAGCAGAAAGAGAGGTGAAACCTATAGACGCTATCACAAATGTCTATCTGATAGACATCGGTACTACCAATGCTATATTGCCAGTTGTGTAATTTACTCTTATTTGATCGTTTCATTTACCAGTTAAGAACCCAAATCATATAAGTGTTATGATGGAAGAACTATAACTTGCAATTCAATTAACTCTGCAATACGATAACAAGCAAAGCGAATCATTTCATTTCGATTTAATCTTTAATTATATATACTTAAACGATGTAAGCCCAAAACAAACGTTTTTTCTATATCTGTCTTTTGAGCAAATTAGTTATACGCAAAACCAAACCGTATTTACATAAATGTATACAAAACAAATCGTATATTTTCATTGGTTTGAAATAAATACATAAAAC
