## Supplementary Figure S1 for "microRNAs affecting development of body pigmentation in adult Drosophila melanogaster"

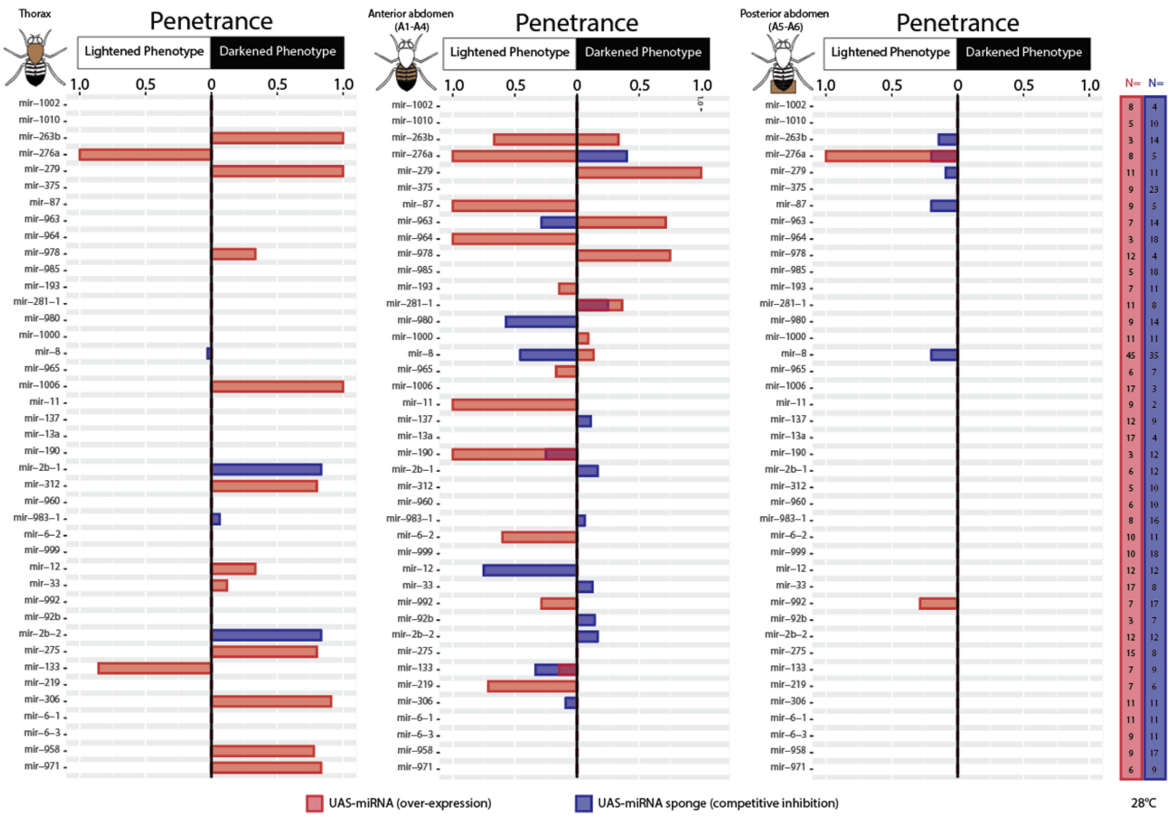


**Supplementary Figure S1. Pigmentation phenotypes caused by overexpression and competitive inhibition of miRNAs in males (at 28ºC).** From left to right, the plots show the penetrance of “lightened” or “darkened” pigmentation phenotypes observed in the thorax, anterior abdomen, and posterior abdomen of male flies. Penetrance ranges from 0 (no flies observed display plotted phenotype) to 1 (all flies observed display plotted phenotype). Penetrance of lightened phenotypes is plotted extending to the left from the “0” axis, while penetrance of darkened phenotypes is plotted extending to the right. Overexpression data is represented by red bars, whereas competitive inhibition data is represented by blue bars. The number of flies scored for each genotype is listed on the right of the figure under “N=” and is color-coded in the same manner as penetrance data (overexpression = red; competitive inhibition = blue). The competitive inhibition data displayed are from flies reared at 28ºC. All lines and genotypes are the same as in **Figure 3**.
