## Supplementary Figure S4 for "microRNAs affecting development of body pigmentation in adult Drosophila melanogaster"

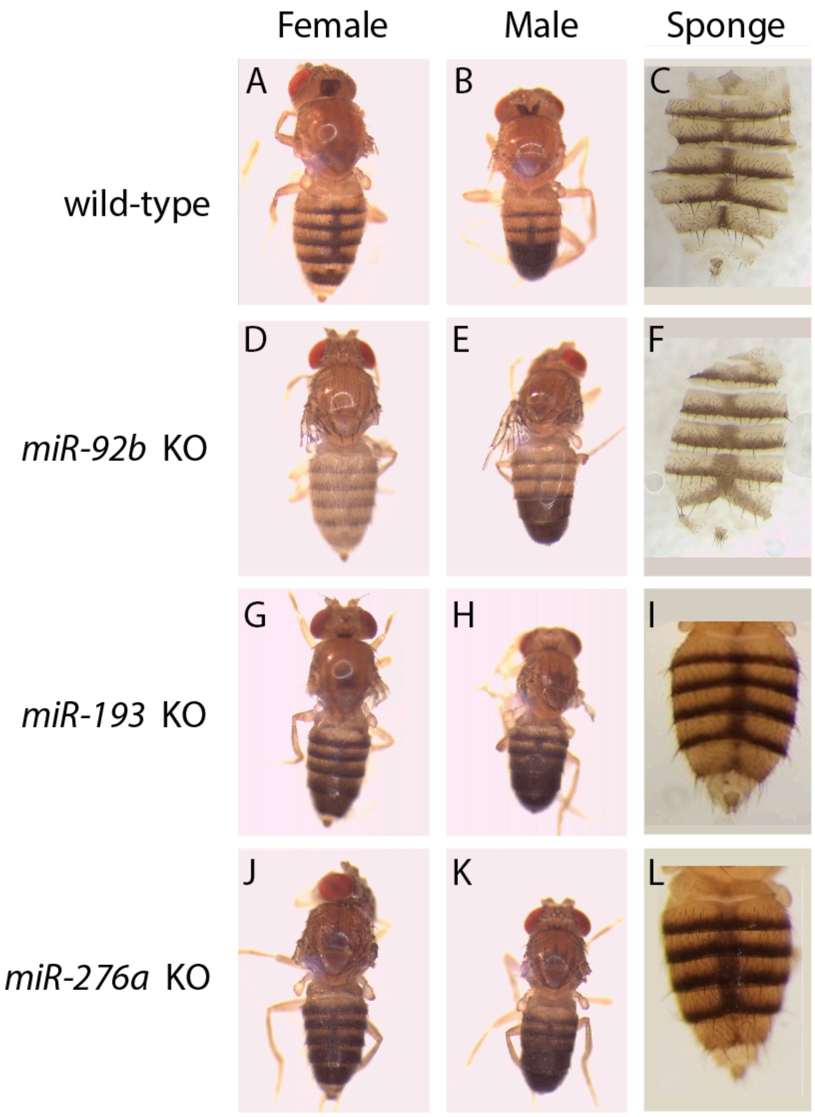


**Supplementary Figure S4**. **Deletion of *miR-92b*, *miR-193,* or *miR-276a* alters body pigmentation.** Three to five day old male (B,E,H,K) and female (A,C,D,F,G,I,J,L) *D. melanogaster* (with dissected dorsal abdominal cuticles in C and F) are shown for genotypes with wild-type pigmentation (*pnr-Gal4* alone in A and B, *pnr-Gal4;UAS-scrambled-sponge* in C), as well as genotypes homozygous for knockout (KO) alleles of *miR-92b* (D,E), *miR-193* (G,H), and *miR-276a* (J,K), or genotypes using *pnr-Gal4* to express a sponge for *miR-92b* (F), *miR-193* (I), or *miR-276a* (L). Panel C is reproduced from Figure 4F (left) and panel F is reproduced from Figure 4F (right).

Panel I is not shown in the main text of this manuscript and is a sibling of the fly shown in Tian et al. (2024) Figure 4B (middle). Panel L is reproduced from Figure 4H (right). For *miR-276a, pnr-Gal4*-driven expression of the sponge showed darker pigmentation in abdominal segments A2-A5 but lighter pigmentation in A6 (L). In flies homozygous for the *miR-276a* knockout allele, all abdominal segments appeared darker (J). For *miR-193*, *pnr-Gal4*-driven expression of the sponge showed lighter pigmentation in abdominal segment A6, with no consistent effect reported for A2-A5 (I). In the knockout allele, all abdominal segments looked darker (G). Finally, for *miR-92a*, expression of the sponge was described as darkening pigmentation in all abdominal segments because it increased the area with darker pigmentation (F). However, the shade of this expanded pigmentation (F) is lighter than the abdominal stripes in the control fly (C), which is similar to the overall lighter pigmentation seen in flies homozygous for the *miR-92b* knockout allele (D). The differences in effects between sponge expression and knockout alleles may be due to the competitive inhibition sponges only reducing miRNA activity in the *pnr-Gal4* expression domain, which is a subset of cells during a subset of development, whereas the flies homozygous for the knockout allele lack the miRNA in all tissues at all developmental stages.
