## Supplementary Figure S5 for "microRNAs affecting development of body pigmentation in adult Drosophila melanogaster"

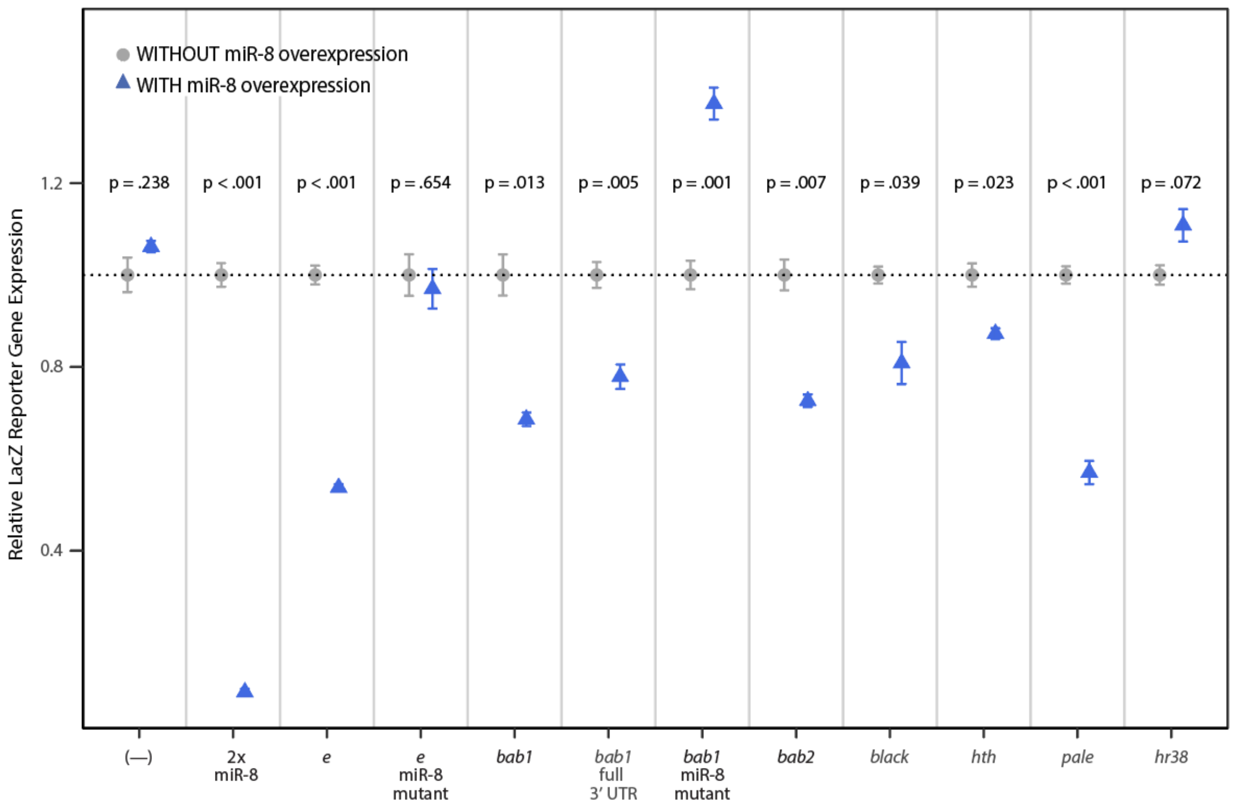


**Supplementary Figure S5. *miR-8* can repress expression of genes via 3' UTR sequences from pigmentation genes.** For each reporter gene, the mean relative LacZ expression (measured as Beta-galactosidase activity/Luciferase activity in S2 cells in each replicate, normalized to the mean expression of that reporter gene without *miR-8* overexpression) is shown. Error bars show the standard error of the mean based on the three replicate measures of each reporter gene in each condition, all collected in parallel on the same day. Data from samples with *miR-8* overexpression are shown in blue; data from samples without *miR*-8 expression are shown in gray. P-values shown for each reporter gene are from t-tests comparing the relative LacZ reporter gene expression between samples with and without *miR-8* overexpression**.** The X-axis labels describe the 3' UTR sequences carried by each LacZ reporter gene. "(—)" = a negative control, no added 3' UTR sequence. "2x miR-8" = a positive control, 3' UTR composed of two perfect complements to the mature *miR-8* sequence. "e" = full *ebony* 3' UTR. "e miR-8 mutant" = same as "e" construct, but with the predicted *miR-8* binding site mutated. “bab1” = a 683 bp region of the *bab1* 3’ UTR sequence containing the predicted *miR-8* binding site. “Bab1 full 3’ UTR”  = full 3’ UTR sequence from *bab1*. “bab1 miR-8 mutant” = same as “bab1” construct, but with the predicted *miR-8* binding site mutated. “bab2” -a 529 bp region of the *bab2* 3’ UTR containing the predicted *miR-8* binding site. “black” - full *black* 3’ UTR. “hth” - full *hth* 3’ UTR.  “pale” - full *pale* 3’ UTR. “hr38” - full *Hr38* 3’ UTR.
